# Cell envelope remodeling requires high concentrations of biotin during *Mycobacterium abscessus* model lung infection

**DOI:** 10.1101/2022.07.20.500843

**Authors:** Mark R. Sullivan, Kerry McGowen, Qiang Liu, Chidiebere Akusobi, David C. Young, Jacob A. Mayfield, Sahadevan Raman, Ian D. Wolf, D. Branch Moody, Courtney C. Aldrich, Alexander Muir, Eric J. Rubin

**Author notes:** Correspondence, Mailing address: 665 Huntington Ave. Building 1, Room 811, Boston, Massachusetts 02115.

## Abstract

*Mycobacterium abscessus* is an emerging pathogen resistant to most frontline antibiotics. *M. abscessus* causes lung infection, predominantly in patients with lung disease or structural abnormalities. To interrogate mechanisms required for *M. abscessus* survival in the lung, we developed a lung infection model using air-liquid interface culture and performed a screen to identify differentially required genes. In the lung model, synthesis of the cofactor biotin is required due to increased intracellular biotin demand, and pharmacological inhibition of biotin synthesis halts *M. abscessus* proliferation. Increased quantities of biotin are required to sustain fatty acid remodeling that serves to increase cell envelope fluidity, which in turn promotes *M. abscessus* survival in the alkaline lung environment. Together, these results indicate that biotin-dependent fatty acid remodeling plays a critical role in pathogenic adaptation to the lung niche and suggests that biotin synthesis and fatty acid metabolism are therapeutic targets for treatment of *M. abscessus* infection.

## Introduction

*Mycobacterium abscessus* is a pathogenic bacterium that has produced an increasing number of human infections over the last two decades^1–3^. Unlike the related organism *Mycobacterium tuberculosis*, *M. abscessus* is not a professional pathogen; it is widespread in the environment and can exist as a free-living bacterium in soil and water. *M. abscessus* can productively infect a large range of organisms, from amoebae to fish to mammals, including humans^4–6^. Though *M. abscessus* can produce systemic infection or localized infection at wound sites, the majority of human disease caused by *M. abscessus* is in the lung^7,8^.

*M. abscessus* predominantly infects individuals with bronchiectasis, chronic obstructive pulmonary disease (COPD), or cystic fibrosis (CF), diverse conditions which share the feature of structurally altered lung parenchyma^9^. During lung infection, *M. abscessus* displays characteristics more akin to opportunistic lung pathogens than to *M. tuberculosis*; for instance, *M. abscessus* resides primarily within the lumen of airways rather than within phagocytic host cells^10,11^, and the majority of end stage *M. abscessus* patients do not display granuloma formation^12^. Thus, *M. abscessus* may represent an intermediate stage in the evolution of an environmental bacterium to a professional pathogen^13,14^.

To survive in the pulmonary milieu, *M. abscessus* must adapt to the conditions within the airways. This environment presents unique challenges; among these obstacles are biophysical stress induced by the presence of mucus^15^, alkaline pH that is higher than most niches in the human body^16,17^, and the requirement to grow in the presence of relatively high oxygen tension at the interface of air and liquid present on the lung surface. The genetic requirements for *M. abscessus* survival in this setting might illuminate the biology that enables environmental bacteria to transition to a pathogenic lifestyle. Furthermore, those constraints may also suggest therapeutic approaches to treat *M. abscessus* infections.

Alternative therapeutic approaches to treat *M. abscessus* are needed, as it is intrinsically resistant to a wide range of antibiotics, and treatment outcomes for individuals with *M. abscessus* infection are poor. Antibiotic treatment produces a cure in only 30-50% of cases, and many patients require surgical resection of lung tissue to control the infection^9,12, 18–23^. Drugs that successfully treat *M. tuberculosis* frequently fail to eradicate *M. abscessus*, and *in vitro M. abscessus* drug susceptibility correlates poorly to antibiotic efficacy in patients^24^. Though *M. abscessus* gene essentiality has been cataloged under standard laboratory conditions^25,26^, environmental context can substantially affect gene essentiality and antibiotic susceptibility^27,28^. Thus, examining genes required for *M. abscessus* infection in a system that models the lung environment may highlight new pathways to target with antibiotics. Many studies have scrutinized how *M. abscessus* behaves in phagocytic cells such as macrophages^29–35^, and a variety of animal models have been developed for *M. abscessus* infection^36–43^; however, these systems often represent systemic or invasive disease, and typically do not recapitulate the extracellular, luminal niche occupied by *M. abscessus* during human lung infection. Tissue culture systems represent an alternative approach to model lung infection, and can be successfully applied to *M. abscessus*^44–47^. Thus, to isolate the role of the lung environment in shaping *M. abscessus* gene essentiality, we have adapted an air-liquid interface tissue culture model for *M. abscessus* infection and used genome-wide saturating transposon mutagenesis and sequencing (TnSeq) to identify genes essential for survival in the lung environment.

## Results

### Development of an air-liquid interface culture system for *M. abscessus* infection

To both recapitulate the environmental conditions present in the lung and preserve tractability to enable genetic screening, we developed an air-liquid interface culture model (Figure 1A) using an immortalized bronchial epithelial cell line, NuLi-1^48^. As previously reported^48^, NuLi-1 cells grown at an air-liquid interface partially differentiate and display features characteristic of bronchial epithelial cells, including epithelial morphology (Figure 1B), partial cilia formation (Supplemental Figure 1A), mucin production (Supplemental Figure 1B), and reduced permeability to small molecules (Supplemental Figure 1C). To monitor whether *M. abscessus* is capable of productively colonizing these lung cell cultures, bacterial luciferase^49^ was constitutively expressed in the *M. abscessus* type strain (ATCC 19977). Luciferase activity correlates well with viable cell numbers in mycobacteria^49^, so *M. abscessus* growth could be observed over time in lung cultures. *M. abscessus* was added apically to lung culture and allowed to attach for 4 hours. Then, liquid was aspirated off of the apical surface to restore the air-liquid interface. A consistent fraction of the bacteria was retained after aspiration (Supplemental Figure 1C), and luminescence was tracked post-aspiration. Infection of lung cell cultures with *M. abscessus* resulted in continuous growth until the cultures reached saturation. The time that cultures took to reach saturation was dependent on the multiplicity of infection (MOI) used for infection (Figure 1D) and was similar for a distantly related clinical isolate of *M. abscessus*, T35 (Supplemental Figure 1E). Infection for 48 hours starting at an MOI of 1 allowed for continuous growth without causing detectable damage to the lung cells as measured by release of the intracellular enzyme lactate dehydrogenase (Figure 1D, Supplemental Figure 1F). As a result, this condition was chosen for all subsequent experiments.

**Figure 1.**
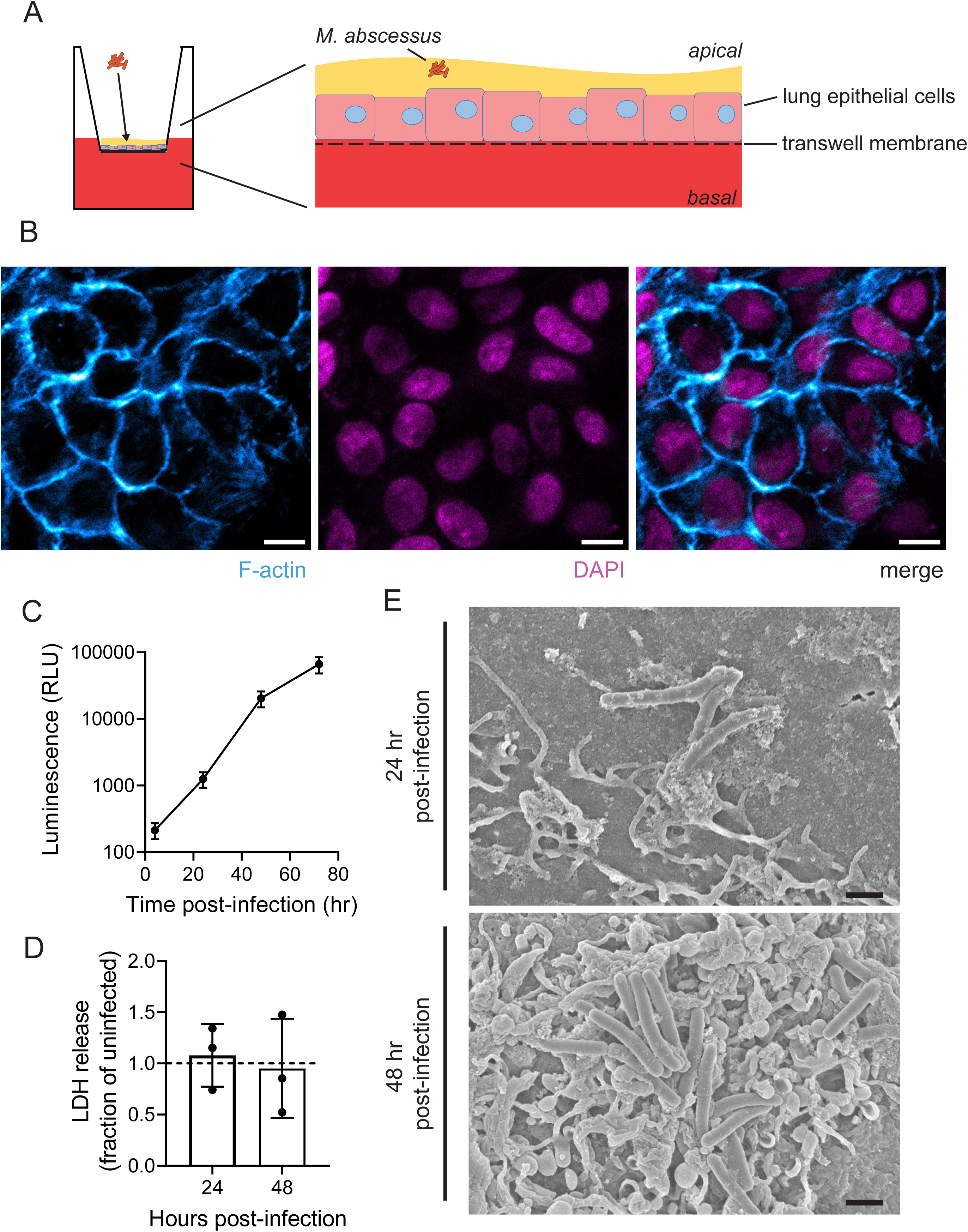
Air-liquid interface culture model for *M. abscessus* lung infection. **(A)** Schematic of air-liquid interface *M. abscessus* culture. **(B)** Confocal microscope images of NuLi-1 lung epithelial cells stained for F-actin and with DAPI to highlight nuclei. Images were obtained at 63X magnification. Scale bar = 10 µm. **(C)** Luminescence measurement of *M. abscessus* expressing bacterial luciferase infected at a multiplicity of infection = 1 on the apical surface of lung epithelial cells. n=3 biological replicates. Data are presented as mean +/− SD. **(D)** Lactate dehydrogenase (LDH) release from lung epithelial cells at 24 and 48 hr post-infection. LDH release is normalized to uninfected control cells. n=3 biological replicates. Data are presented as individual values along with mean +/− SD. **(E)** Scanning electron microscope images of apical surface of lung infection model at 24 and 48 hr post-infection. Images were obtained at 11000X magnification. Scale bar = 1 µm.

In human lung, *M. abscessus* forms aggregates on the surface of epithelial cells^10,11^, a behavior that has been observed in other models for *M. abscessus* lung infection^44^. To examine whether our air-liquid interface model recapitulates this physiological mode of *M. abscessus* growth, infected cultures were observed by scanning electron microscopy (SEM) at 24 and 48 hours after infection. At both time points, *M. abscessus* was visible growing on the surface of the lung cell layer (Figure 1E). To test whether the lung epithelial cells were also phagocytosing *M. abscessus*, a fluorescent strain of *M. abscessus* expressing mScarlet^50^ was generated and used to infect lung cultures. After washing away all surface-attached bacteria, fluorescence microscopy revealed that phagocytosed *M. abscessus* were present (Supplemental Figure 1G-H), but very rare in comparison to the fraction of cells growing on the surface of the epithelial layer (Figure 1E). Together, these data suggest that this model can effectively recapitulate clinical characteristics of *M. abscessus* lung infection.

### Genetic screening in lung infection model

To interrogate the genetic requirements for survival and growth in an environment mimicking the lung, we carried out a genetic screen in the lung infection model using TnSeq^51^. To identify genes that might be relevant in the context of infection, we utilized a clinical isolate of *M. abscessus*, T35, that has not been passaged in culture as the *M. abscessus* type strain has. Gene essentiality was compared among 3 conditions: the input library grown on standard agar plates, *M. abscessus* grown in the lung infection model, and *M. abscessus* cultured directly in the tissue culture medium used for the lung model. This tissue culture medium is relatively close in composition to human serum (Supplemental File 1) and represents a more physiological environment than mycobacterial culture medium^28,52^. Further, tissue culture medium supports proliferation at a similar rate as the lung infection model (Supplemental Figure 2A), which allows direct comparisons between screen conditions. We reasoned that comparison of gene essentiality among these three conditions might both illuminate genes important for growth in more physiological nutrient conditions and genes that are specifically made essential by growth on the surface of lung cells.

### Biotin biosynthetic enzymes are required for growth in lung infection model

Growth in the lung infection model renders 237 genes significantly depleted for transposon insertions compared to the input library (Figure 2A, Supplemental File 2, Supplementary Table 1). To isolate the genes most critical for lung infection, we focused primarily on genes that are further required in the lung infection model beyond their need in tissue culture medium. From this more stringent comparison, the two genes most differentially required in the presence of lung cells are both members of the biotin biosynthetic pathway (Figure 2B, Supplemental Figure 2B)^53^, suggesting that growth in the lung infection model imposes an increased requirement for biotin synthesis. Further, all of the members of the biotin biosynthetic pathway display greatly reduced insertion counts in the lung infection model (Figure 2C), consistent with an increased need for endogenous biotin synthesis. Of note, genes in the biotin synthesis pathway are significantly more required in tissue culture medium alone than on agar plates, but the presence of lung epithelial cells further increases the requirement for biotin biosynthetic genes (Figure 2C). Indeed, deletion of *bioA* (*MAB_2688c*) (Supplemental Figure 2C-E) impedes growth both in the lung infection model and in tissue culture medium (Figure 2D). This growth defect is a result of biotin insufficiency, as high levels of exogenous biotin or re-expression of BioA can rescue growth of the Δ*bioA* strain (Figure 2D).

**Figure 2.**
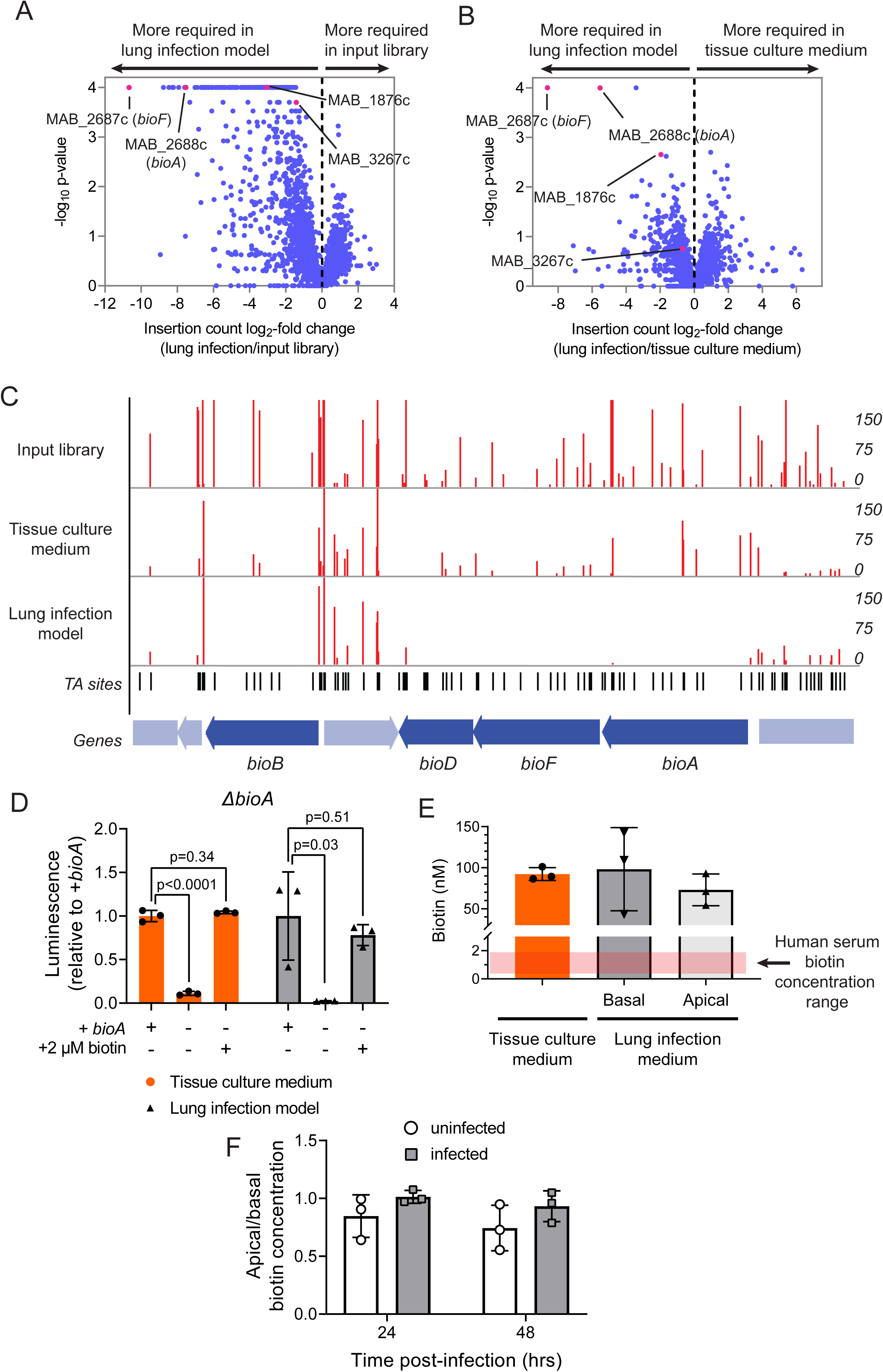
Biotin synthesis is required in culture media and lung infection model despite presence of biotin. **(A)** Log_2_-fold ratio of transposon insertion counts plotted against significance in **(A)** the lung infection model versus the input library and **(B)** the lung infection model versus tissue culture medium. p-values derived from permutation test. **(C)** Insertion counts for indicated genes in a representative replicate of the input library, tissue culture medium, or lung infection model. Insertion counts are normalized to the local maximum. **(D)** Luminescence of *ΔbioA M. abscessus* ATCC19977 with (*bioA* +) or without (*bioA*−) BioA expression genetically rescued after 48 hr in either tissue culture medium or in the lung infection model. Culture medium contained either no supplemental biotin or supplementation of 2 µM biotin. Data are presented as individual values along with mean +/− SD. n = 3 biological replicates. p-values derived from unpaired, two-tailed t-test. **(E)** Biotin concentration measured by enzyme-linked immunosorbent assay (ELISA) in either tissue culture medium or liquid taken from the apical or basal compartments of a mature air-liquid interface culture after 48 hours. Red bar represents the range of reported human serum biotin concentrations^54^. Data are presented as individual values along with mean +/− SD. n = 3 biological replicates. **(F)** Ratio of biotin concentration measured by ELISA in the apical and basal compartments of the air-liquid interface culture model. Medium was sampled from either uninfected or infected air-liquid interface cultures at 24 and 48 hours after initiation of infection or mock infection. Data are presented as individual values along with mean +/− SD. n = 3 biological replicates.

### Lung infection model imposes increased demand for biotin synthesis despite biotin in medium

Biotin is present in the lung culture medium at levels 20-100 times greater than in human serum^54^ (Figure 2E), though at lower levels than in standard mycobacterial medium (Supplemental File 1). Free biotin levels available in the lung infection model do not differ from those present in tissue culture medium alone (Figure 2E), nor do lung cells selectively deplete biotin on the apical surface compared to basal medium (Figure 2F), suggesting that altered biotin availability does not fully explain the differential requirement for biotin synthetic enzymes. Thus, we questioned whether growth in more physiological environments might cause bacteria to have a higher absolute demand for biotin.

To evaluate whether growth in tissue culture medium increases demand for biotin, we cultured the *ΔbioA M. abscessus* strain in increasing concentrations of exogenous biotin to determine the minimal amount of biotin required to sustain proliferation. When grown in tissue culture medium, *ΔbioA M. abscessus* requires a higher concentration of exogenous biotin to support growth than when cultured in mycobacterial medium (Figure 3A). This phenomenon holds true in tissue culture medium lacking all supplements and protein components (Figure 3A), suggesting that there are not factors present sequestering biotin in tissue culture medium. Similarly, conditioned media taken from the lung infection model that was dialyzed to retain protein components while refreshing small molecule metabolites does not impose an altered demand for biotin synthesis (Supplemental Figure 3A), arguing that lung cells do not produce protein factors that sequester biotin.

**Figure 3.**
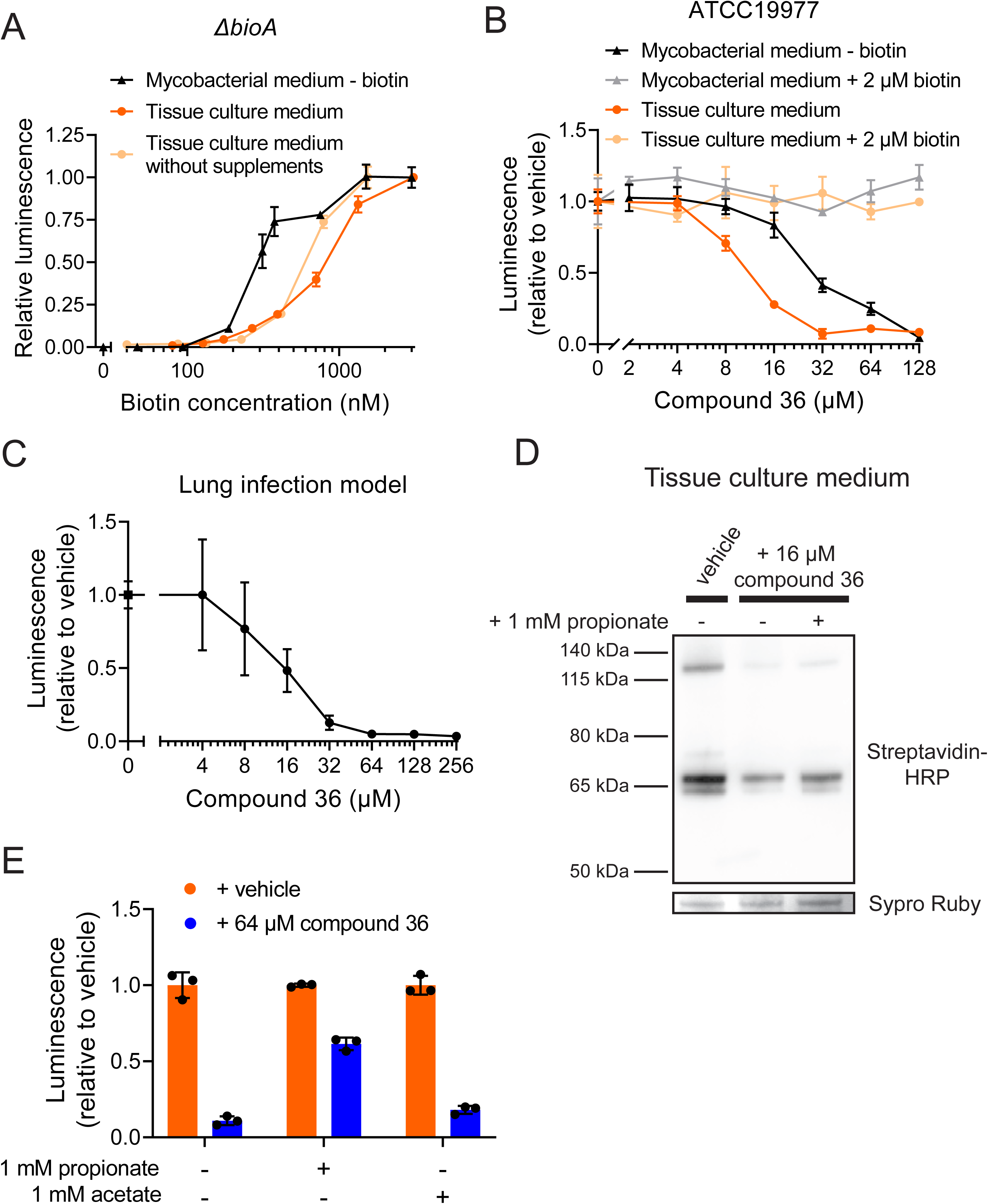
Physiological environments impose demand for biotin synthesis. **(A)** Luminescence of *ΔbioA M. abscessus* ATCC19977 grown in the indicated medium for 48 hr with the specified final concentrations of biotin in the medium. Values are normalized within each medium to 3 µM biotin. **(B)** Luminescence of *M. abscessus* ATCC19977 grown in the indicated medium for 48 hr with the specified final concentrations of the BioA inhibitor compound 36. Values are normalized within each medium to the vehicle treated condition. **(C)** Luminescence of *M. abscessus* ATCC19977 grown in the lung infection model for 48 hr with the specified final concentrations of compound 36. Values are normalized to the vehicle treated condition. **(D)** Western blot for total biotinylated protein in *M. abscessus* ATCC19977 grown in tissue culture medium with either vehicle or 16 µM compound 36 along with the indicated supplementation of propionate. A representative band of SYPRO Ruby staining for total protein is displayed for each condition. **(E)** Luminescence of *M. abscessus* ATCC19977 grown in tissue culture medium for 48 hr with either vehicle or 64 µM compound 36 along with the indicated supplementation of propionate or acetate. Values are normalized within each condition to vehicle-treated. For all graphs, data are presented as mean +/− SD. n = 3 biological replicates.

To orthogonally examine whether *M. abscessus* requires more biotin in tissue culture medium, we sought to determine whether *M. abscessus* was more susceptible to inhibition of biotin synthesis. Inhibitors of biotin synthesis have been developed for use in *M. tuberculosis*^55–58^, and we found that an *M. tuberculosis* Rv1568 (BioA) inhibitor, compound 36 (PubChem CID: 137348519)^57^, is effective at inhibiting *M. abscessus* growth in the absence of biotin (Figure 3B). Compound 36 does not inhibit growth in the presence of biotin (Figure 3B), indicating that its anti-proliferative effects are specifically caused by biotin synthesis inhibition. BioA inhibition impedes proliferation more effectively in tissue culture medium than in mycobacterial medium (Figure 3B), consistent with an increased demand for biotin in the more physiological medium. Further, BioA inhibition by compound 36 prevents growth in an assortment of *M. abscessus* clinical isolates (Supplemental Figure 3B) and is active in the lung infection model (Figure 3C, Supplemental Figure 3C) without detectable toxicity to the lung cells (Supplemental Figure 3D), which suggests that BioA inhibition may represent a therapeutic strategy for treating *M. abscessus* infection.

### Biotin synthesis inhibition is rescued by propionate metabolism

An increased demand for biotin synthesis suggests a larger requirement for biotin-utilizing enzymes. *M. abscessus* possesses several biotin-dependent enzymes^59,60^, and these proteins broadly display diminished biotinylation upon biotin synthesis inhibition (Figure 3D). Two biotin-dependent enzymes are significantly more required in the lung infection model than in the input library: a pyruvate carboxylase, *MAB_3267c*, and an acetyl-CoA/propionyl-CoA carboxylase, *MAB_1876c*. Pyruvate carboxylase is essential for induction of biotin synthesis in mycobacteria^61^, so we instead focused on whether an activity catalyzed by the acetyl-CoA/propionyl-CoA carboxylase gene is more critical in physiological environments than in mycobacterial medium. We posited that if metabolism of acetyl-CoA or propionyl-CoA was selectively more important for growth in physiological environments, we might be able to rescue partial biotin synthesis inhibition by adding excess acetate or propionate to drive forward those metabolic pathways. Indeed, propionate rescues growth upon BioA inhibition, while acetate fails to rescue (Figure 3E). Propionate rescues growth without increasing protein biotinylation (Figure 3D), suggesting that propionate acts downstream of biotin and allows cells to proliferate despite low biotin levels. Further, cholesterol, a biologically relevant source of propionate for mycobacteria^62^, rescues biotin synthesis inhibition (Supplemental Figure 3E). Together, these results suggest that downstream utilization of propionate partly protects *M. abscessus* against biotin synthesis inhibition.

### Physiological medium imposes altered demands for fatty acid synthesis

Propionate has three major fates in mycobacteria; it can be used to generate methyl-branched fatty acids, synthesize odd-chain fatty acids, or can be recycled back into the tricarboxylic acid (TCA) cycle through either the methylcitrate cycle or through methylmalonyl-CoA epimerase and mutase^63^. Given that all of the genes required for recycling of propionyl-CoA into the TCA cycle are non-essential in all tested conditions (Supplemental Table 2), we focused on whether growth in physiological environments imposes altered demands for fatty acid synthesis. Using gas chromatography/mass spectrometry (GC/MS), we determined that the profile of fatty acids in *M. abscessus* cultured in tissue culture medium differs from that of cells grown in mycobacterial medium (Figure 4A, Supplemental File 3). Strikingly, growth in tissue culture medium increases the number of branched, unsaturated, and odd-chain fatty acids with concomitant decreases in most even-chain saturated fatty acid species.

**Figure 4.**
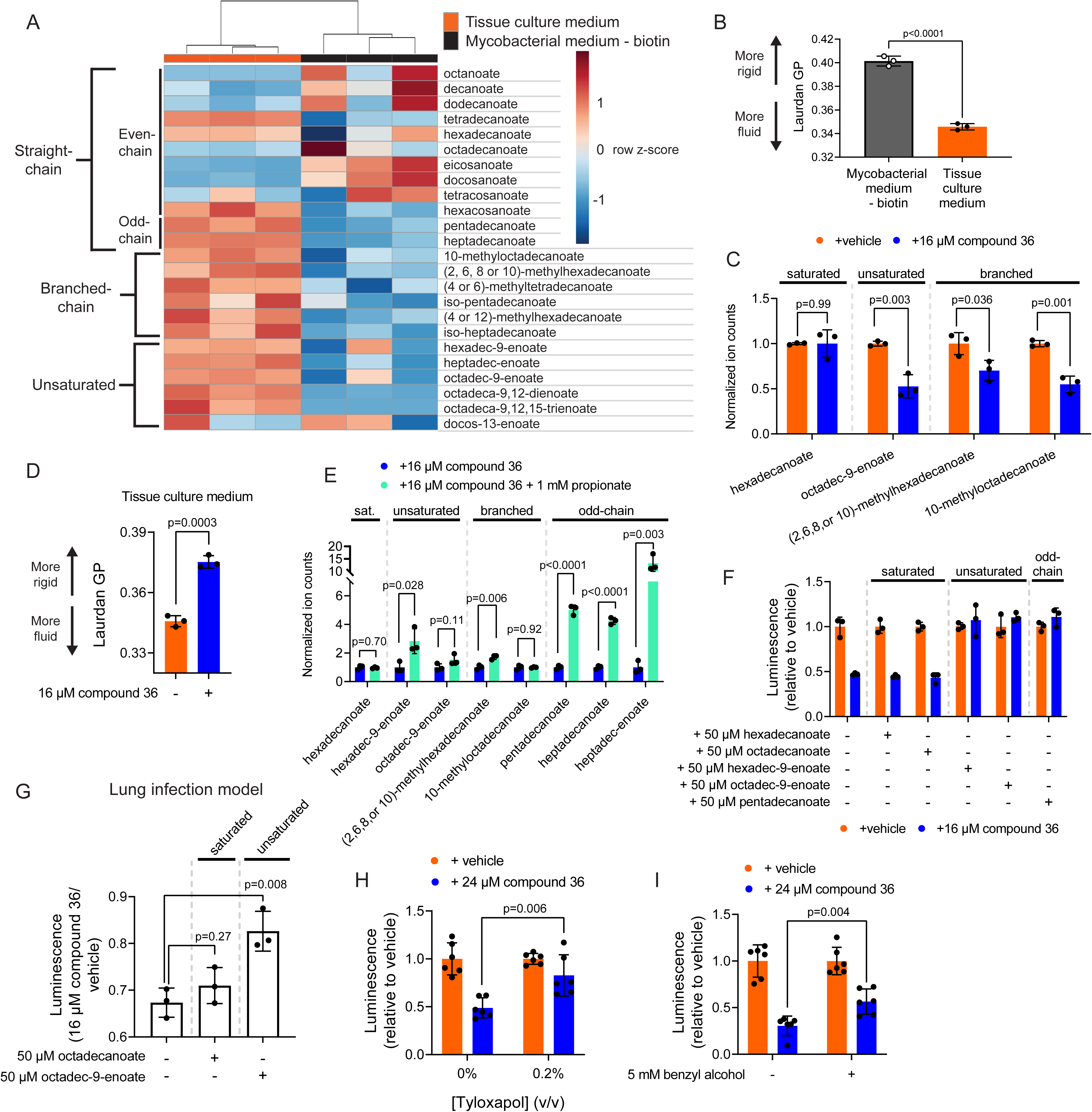
Biotin is required to support fatty acid remodeling that sustains envelope fluidity. **(A)** Heatmap depicting relative abundance of 24 fatty acid species measured by GC/MS in *M. abscessus* ATCC19977 grown 48 hr in either tissue culture medium or mycobacterial medium - biotin. Samples are hierarchically clustered, while fatty acid species are ordered by their class and are not clustered. **(B)** Laurdan generalized polarization (GP) for *M. abscessus* ATCC19977 grown 48 hr in either tissue culture medium or mycobacterial medium - biotin. Data are presented as individual values along with mean +/− SD. n=3 biological replicates. p-value derived from unpaired, two-tailed t-test. **(C)** GC/MS measurement of the indicated fatty acids in *M. abscessus* ATCC19977 grown 48 hr in tissue culture medium treated with either vehicle or 16 µM compound 36. **(D)** Laurdan generalized polarization (GP) for *M. abscessus* ATCC19977 grown 48 hr in tissue culture medium treated with either vehicle or 16 µM compound 36. **(E)** GC/MS measurement of the indicated fatty acids in *M. abscessus* ATCC19977 grown 48 hr in tissue culture medium treated with 16 µM compound 36 along with either vehicle or 1 mM sodium propionate. **(F)** Luminescence of *M. abscessus* ATCC19977 grown in tissue culture medium for 48 hr treated with either vehicle or 16 µM compound 36 along with the indicated supplementation of fatty acids. Values are normalized within each condition to vehicle-treated. **(G)** Ratio of luminescence of *M. abscessus* ATCC19977 in air-liquid interface lung cultures treated with 16 µM compound 36 compared to vehicle-treated. Basal medium was supplemented with the indicated fatty acids, and infections lasted 48 hr. **(H)** Luminescence of *M. abscessus* ATCC19977 grown in tissue culture medium for 48 hr treated with either vehicle or 24 µM compound 36 along with the indicated concentration of tyloxapol **(I)** or benzyl alcohol **(H)**. Values are normalized within each condition to vehicle-treated. n = 6 biological replicates. For all graphs, data are presented as individual values along with mean +/− SD. n = 3 biological replicates unless otherwise indicated. All p-values derived from unpaired, two-tailed t-tests.

To determine whether this shift in fatty acid composition has biologically relevant consequences, we tested whether the physical characteristics of the cell envelope are different in *M. abscessus* cultured in tissue culture medium compared to mycobacterial medium. Membrane fluidity is known to increase with higher fractional composition of unsaturated and branched fatty acids^64^, so we predicted that fluidity of the cell envelope would increase in *M. abscessus* cultured in tissue culture medium. To assess envelope fluidity, we adapted a previously described assay using laurdan^65^, a dye that displays shifts in the maximum of its fluorescence emission spectrum based on the fluidity of the surrounding membrane^66,67^. Laurdan can be used to probe bulk cell envelope fluidity in *M. abscessus* (Supplemental Figure 4A-B), similar to methods used in other microbes^68,69^. The laurdan generalized polarization (GP) (see Materials and Methods) is a standardized ratio of fluorescence emission intensities that anti-correlates with membrane fluidity^70^. The observed shift towards less saturated fatty acid chains in tissue culture medium is accompanied by an increase in envelope fluidity, which is indicated by a decrease in laurdan GP (Figure 4B). This change in cell envelope physical properties signifies that substantial membrane remodeling has occurred, and suggests that culture medium may alter demand for fatty acid synthesis and, therefore, requirement for biotin.

### Biotin availability supports fatty acid remodeling

To address whether fatty acid composition is impacted by biotin deficiency, fatty acid abundance was measured upon partial inhibition of BioA (Supplemental Figure 4C). A selection of unsaturated and branched fatty acids were depleted by BioA inhibition, with no changes observed to the abundant straight-chain fatty acid, hexadecanoate (Figure 4C). Notably, BioA inhibition results in a decrease in envelope fluidity as indicated by increased laurdan GP (Figure 4D). Together, these results suggest that BioA inhibition results in meaningful membrane remodeling and that biotin availability is required to sustain production of non-straight chain fatty acids that make up a larger fraction of the membrane in physiological environments.

### BioA inhibition is rescued by envelope fluidizing agents

Since propionate rescues BioA inhibition, we hypothesized that propionate supplementation would restore synthesis of fatty acids depleted by BioA inhibition. However, while propionate provided minor rescues to some unsaturated and branched fatty acids, its most notable effect was to dramatically increase synthesis of odd-chain fatty acids (Figure 4E, Supplemental Figure 4D). Odd-chain fatty acids also increase membrane fluidity^71,72^, while bypassing the biotin-dependent propionyl-CoA carboxylase reaction that is required to utilize propionate for methyl-branched fatty acid synthesis (Supplemental Figure 4E). This result suggests that BioA inhibition does not deprive *M. abscessus* of a single critical fatty acid, but instead alters the envelope’s physical properties in a way that can be remedied by various non-straight chain fatty acids. Consistent with this model, supplementation of either unsaturated or odd-chain fatty acids rescues BioA inhibition, while provision of saturated, even-chain fatty acids does not (Figure 4F). Further, exogenous supplementation of the unsaturated fatty acid (9Z)-octadec-9-enoate rescues biotin synthesis inhibition in the lung infection model, while provision of a saturated, even-chain fatty acid, octadecanoate, does not (Figure 4G).

Since fatty acids can be metabolized by *M. abscessus* and potentially produce secondary effects, we sought to determine whether modulating envelope fluidity through non-metabolizable chemical interventions could rescue biotin deprivation. Supplementation of the detergent tyloxapol, which is not metabolized by cells^73^ but alters envelope properties, partially rescues BioA inhibition (Figure 4H). Similarly, addition of benzyl alcohol, a membrane fluidizing agent^74^ that also alters membrane partitioning^75^, allows *M. abscessus* to better tolerate biotin synthesis inhibition (Figure 4I). Together, these results argue that biotin is required in more physiological environments to maintain synthesis of fatty acid species that increase envelope fluidity.

### Anti-proliferative effects of BioA inhibition are not caused by depletion of a specific lipid

Though these results are consistent with a model in which modulation of bulk envelope properties is the primary effect of BioA inhibition, we sought to evaluate the possibility that biotin deficiency also inhibits proliferation by curtailing production of a specific, critical lipid. Given that changes in a single lipid species might not be detectable by bulk methods like fatty acid methyl ester analysis, we characterized the lipid content of *M. abscessus* grown in tissue culture medium using high performance liquid chromatography/mass spectrometry (HPLC/MS)^76,77^. Upon BioA inhibition, the majority of differentially abundant compounds detected by HPLC/MS were directly derived from the BioA inhibitor, compound 36, and only one compound was significantly depleted by BioA inhibition (Supplemental Figure 4F-G). Since propionate is sufficient to rescue *M. abscessus* growth under these conditions, propionate supplementation would be predicted to rescue the abundance of any critical, depleted lipid species that are required for proliferation. However, propionate fails to rescue the compound significantly depleted upon BioA inhibition (Supplemental Figure 4F), suggesting there is not a specific lipid that mediates the anti-proliferative effects of BioA inhibition. Collectively, these results support a model in which BioA inhibition is predominantly deleterious to *M. abscessus* by preventing bulk membrane remodeling required to promote envelope fluidity.

### Alkaline pH imposes demand for biotin and alters fatty acid composition of *M. abscessus*

Given that tissue culture medium imposes an increased demand for biotin to support non-straight chain fatty acid synthesis, we posited that some element of the physiological environment must create a stress that necessitates a change in cell envelope properties. To identify this stress, we added pools of the individual components of mycobacterial medium to tissue culture medium to determine whether any components of mycobacterial medium could rescue biotin synthesis inhibition. We observed a striking correlation between the medium pH and sensitivity to BioA inhibition (Supplemental Figure 5A) that was independent of the nutrients present in each pool. Indeed, lowering the pH of tissue culture medium from 7.8 to 6.8 to match mycobacterial medium decreases sensitivity to BioA inhibition (Figure 5A), without rescuing protein biotinylation (Figure 5B). This suggests that the activity of the BioA inhibitor is not altered by pH, as biotin levels are unchanged. Further, increasing the pH of mycobacterial medium increases sensitivity to BioA inhibition (Figure 5C), indicating that the pH of the medium represents one determinant of biotin synthesis demand.

**Figure 5.**
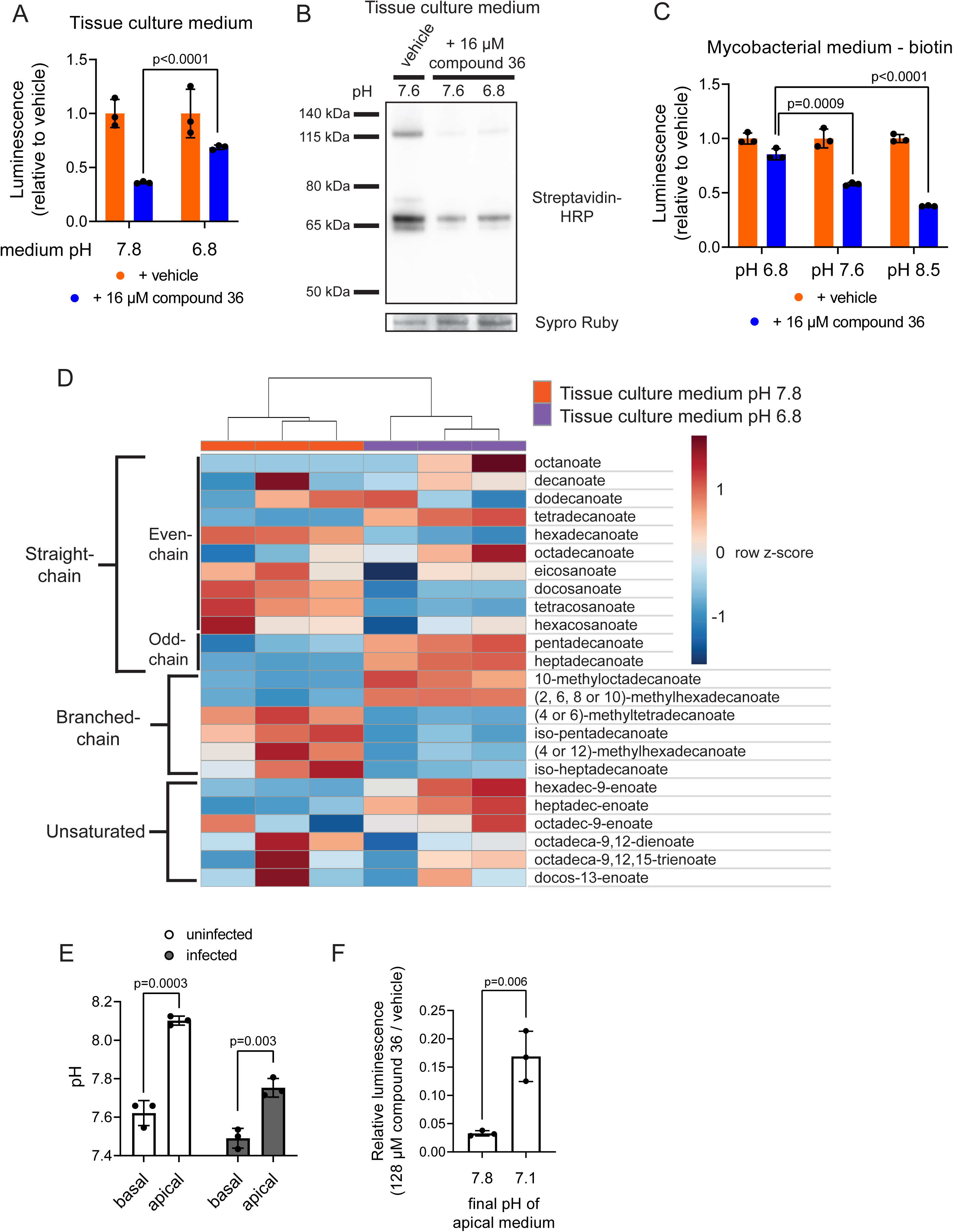
Physiological pH alters fatty acid profile and imposes increased demand for biotin. **(A)** Luminescence of *M. abscessus* ATCC19977 grown in tissue culture medium adjusted to the indicated pH and treated with either vehicle or 16 µM compound 36 for 48 hr. Values are normalized within each condition to vehicle-treated. **(B)** Western blot for total biotinylated protein in *M. abscessus* ATCC19977 grown in tissue culture medium adjusted to the indicated pH and treated with either vehicle or 16 µM compound 36. A representative band of SYPRO Ruby staining for total protein is displayed for each condition. **(C)** Luminescence of *M. abscessus* ATCC19977 grown in mycobacterial medium - biotin adjusted to the indicated pH and treated with either vehicle or 16 µM compound 36 for 48 hr. Values are normalized within each condition to vehicle-treated. **(D)** Heatmap depicting relative abundance of 24 fatty acid species measured by GC/MS in *M. abscessus* ATCC19977 grown 48 hr in tissue culture medium adjusted to the indicated pH. Samples are hierarchically clustered, while fatty acid species are ordered by their class and are not clustered. **(E)** pH of liquid sampled from the basal and apical surfaces of infected or mock infected air-liquid interface lung cultures as measured by phenol red absorbance. **(F)** Ratio of luminescence of *M. abscessus* ATCC19977 in air-liquid interface lung cultures treated with 128 µM compound 36 compared to vehicle-treated after 48 hr infection. Initial basal pH was adjusted to either 7.6 or 6.8, and final apical pH in each condition was determined to be 7.8 and 7.1, respectively. For all graphs, data are presented as individual values along with mean +/− SD. n = 3 biological replicates. All p-values derived from unpaired, two-tailed t-tests.

Increases in environmental pH correlate with increases in branched and unsaturated fatty acid content^78–80^, and acidic stress leads to an altered fatty acid profile^78,79^, suggesting that extracellular pH may influence fatty acid metabolism. To address how alkaline stress might increase demand for biotin, fatty acid abundance was measured in *M. abscessus* grown in tissue culture medium either at pH 7.8 or pH 6.8. Fatty acid composition in *M. abscessus* is altered at pH 6.8 (Figure 5D, Supplemental Figure 5C-D) despite the relatively small shift in pH, and the altered fatty acid profile at pH 6.8 suggests two mechanisms by which lower pH reduces demand for biotin synthesis. First, growth at pH 6.8 leads to decreased levels of many branched fatty acids (Figure 5D), suggesting that *M. abscessus* has a lower demand for their synthesis at pH 6.8. Second, lower pH increases the abundance of several unsaturated fatty acids (Figure 5D), which might partially alleviate the demand for increased non-straight chain fatty acid synthesis.

### Alkaline pH stress is exacerbated by lung cells

Recent studies make clear that the fluid that lines the human airway is regulated to an alkaline pH^16,17^, which might serve as a luminal anti-bacterial mechanism^16,81^. Thus, we questioned whether the increased requirement for biotin synthesis in the lung infection model is driven by heightened alkaline stress generated by the lung cells beyond that present in tissue culture medium. NuLi-1 cells increase the alkalinity of their apical surface, both in monoculture and during *M. abscessus* infection (Figure 5E) to a degree that closely matches the alkalinity of the upper human airway^17^. This suggests that exacerbation of alkaline stress may contribute to the increased biotin synthesis demand in the lung environment. Consistent with this possibility, reducing the pH of the culture medium in the lung infection model partially rescues the growth defect caused by biotin synthesis inhibition (Figure 5F), though the degree of rescue is limited by the continuous alkalinization of the apical medium by the lung cells (Supplemental Figure 5E). Together, these results suggest that the lung environment imposes alkaline stress on *M. abscessus* that necessitates a shift in fatty acid profile that increases demand for biotin.

## Discussion

Biotin synthesis has long been seen as an attractive target for antibiotic therapy^55–58,82–86^, as mammals lack homologous enzymes, and prior work in *M. tuberculosis* has shown that biotin synthesis is essential *in vivo*^85,87^. Most effort towards clinical inhibition of biotin synthesis has been in intracellular pathogens, with the rationale that these organisms will experience biotin starvation caused by sequestration inside phagocytic cells. However, recent work in surface-dwelling lung pathogens has suggested that these organisms may also be susceptible to biotin synthesis inhibition, but that their sensitivity had been overlooked due to poor representation of human biotin levels in mouse models of infection^88^. Similarly, our results suggest that biotin synthesis represents a critical step for bacteria that are growing on the apical surface of the human lung and that biotin synthesis inhibition may represent an effective therapy. Further, given that biotin synthesis inhibition acts to limit growth of extracellular bacteria present in the lung cavity, delivery of biotin synthesis inhibitors, either systemically or via aerosol, may enable delivery of high concentrations of these drugs with limited side effects and could be combined with other *M. abscessus* antibiotics that are delivered through inhalation^24,89–91^.

The apical surface of the lung is known to be a basic environment^16,17^, and the growth of pathogens is hindered by this alkalinity^16^. Proper alkalinization of the apical surface of the lung is impaired in various disease states that predispose individuals to *M. abscessus* infection, notably cystic fibrosis^16^. Given the role of alkalinization in pathogen defense, recent work has focused on artificial alkalinization of the lung surface to augment immune defense against pathogens^92–96^. Potential synergy between alkalinizing treatments and biotin synthesis inhibition might represent a method for eradicating opportunistic pathogens in vulnerable individuals. Of note, alkaline pH appears to inhibit a wide range of clinically important lung pathogens^16^, many of which synthesize biotin *de novo* and might be sensitive to pharmacological biotin deprivation ^88^.

Despite the evidence that elevated lung pH plays an important role in pathogen defense, most consequences of dysfunctionally low pH have been linked with impaired host cell function^93^ rather than with direct effects on bacteria. Some work has suggested that alkaline environments directly hinder bacterial growth through high bicarbonate concentrations, rather than through pH changes themselves^97,98^, and extracellular bicarbonate concentration has a marked impact on the bacterial transmembrane pH gradient^99^. Bicarbonate abundance does not appear to be the sole causative factor underlying alkaline stress on *M. abscessus* given that high pH is still detrimental to growth in mycobacterial medium without biotin, which lacks bicarbonate (Figure 5C); however, bicarbonate levels are likely increased in the alkalinized lung lumen, as well as in alkalinized culture medium^100^, and thus may play an additional role in limiting *M. abscessus* proliferation.

Regardless of the relative contributions of pH and bicarbonate in the lung environment, we find that the stress imposed upon *M. abscessus* by high pH can be counteracted by fatty acid remodeling that increases envelope fluidity, and that exogenous addition of fluidizing agents can also alleviate this stress. Of note, several biosynthetically unique classes of fatty acids counteract the excess demand for biotin and fatty acid synthesis, emphasizing that the increased biotin requirement observed in the alkaline lung environment appears to be driven by a need to adjust the physical properties of the envelope, rather than by a need for a specific lipid species. This altered requirement for envelope fluidity suggests a number of possible mechanisms by which alkaline environments might be deleterious to bacterial growth. One compelling candidate that might be influenced by both envelope properties and extracellular pH is the activity and localization of membrane proteins. Membrane properties are critical to support proper folding of membrane proteins, as membrane tension created by lipids with non-linear fatty acid chains or conical head groups tends to promote appropriate folding and insertion of proteins into the membrane^101^. Further, membrane fluidity is a critical determinant of the essential process of membrane partitioning displayed by many bacterial species^69^, including mycobacteria^75^, and alterations in partitioning may play a central regulatory role in membrane protein function^69^. In addition, membrane composition is coupled to protein oligomerization^102^, and fluid membranes are required for the activity of respiratory complexes that rely upon diffusion of factors through the membrane^103^, providing several avenues by which membrane composition might impact protein function.

Similarly, alkaline pH can affect membrane proteins in many ways. Low extracellular proton concentration could impair proton-gradient dependent proteins, which constitute a large fraction of cell-surface proteins. Additionally, direct exposure to high pH might cause extracellular domains of proteins to unfold and function poorly, which could impose a higher demand on membrane properties to maintain those proteins in a folded state. Further exploration of which of these mechanisms are relevant might suggest additional therapeutic synergies with both biotin synthesis inhibition and alkaline pH.

Together, these results suggest heretofore undescribed connections between the alkaline environment of the lung and biotin-dependent fatty acid remodeling that serves to preserve envelope fluidity. Future work to examine these connections may further illuminate the mechanisms by which fatty acid remodeling is used by pathogens to respond to environmental stresses like alkalinity and suggest additional therapeutic avenues for treating apical lung pathogens.

## Acknowledgements

We acknowledge all of the members of the Rubin and Fortune labs for input and advice on the manuscript, and we thank Jin-Ah Park and Chimwemwe Mwase for assistance with air-liquid interface cultures. We thank the Biopolymers Facility at Harvard Medical School for sequencing and the Microscopy Resources on the North Quad (MicRoN) core at Harvard Medical School for assistance with microscopy. Electron microscopy imaging was performed in the HMS Electron Microscopy Facility. M.R.S. is a Merck Fellow of the Damon Runyon Cancer Research Foundation, DRG-2415-20. E.J.R. was supported by a Dean’s Innovation Award from Harvard Medical School, and by NIH/NIAID under award number R21AI156772. A.M. acknowledges support from the Ludwig Center for Metastasis. D.B.M acknowledges R01 AI049313 and U19 AI162584.

## Author Contributions

Conceptualization, M.R.S., E.J.R.; Methodology, M.R.S., C.A.; Formal Analysis, M.R.S., D.C.Y., J.A.M.; Investigation, M.R.S., K.M., I.D.W., D.C.Y., S.R., A.M.; Software, J.A.M.; Resources, Q.L., C.C.A.; Visualization, M.R.S., D.C.Y., J.A.M.; Writing – Original Draft, M.R.S.; Writing – Second Draft, Review & Editing, M.R.S., K.M., C.A., D.C.Y., J.A.M., A.M., D.B.M., E.J.R.; Funding Acquisition, M.R.S., E.J.R.; Supervision, E.J.R., C.C.A., D.B.M.

## Declaration of Interests

D.B.M consults with Pfizer and EnaraBio. The other authors declare no competing interests.

## Materials Availability

All reagents generated in this study are available upon request from the corresponding author.

## Data Availability

All relevant data generated in this study are present within the manuscript and Supplemental Information, with the following exceptions. Whole genome sequencing data for strain T35 are available on SRA (https://www.ncbi.nlm.nih.gov/sra) under project number PRJNA840944, accession number SAMN28571509. Raw TnSeq data are available on SRA, and the project and accession numbers will be listed prior to publication.

## Methods

### Strains

All experiments were performed in the *M. abscessus abscessus* type strain (ATCC 19977) unless otherwise indicated. Clinical isolates of *M. abscessus* were isolated from patients in Taiwan. Strain T35 was isolated from a surgical wound site, T37 was isolated from a lung biopsy, and strains T40 and Bamboo^104^ were isolated from sputum. All plasmid construction was performed in DH5α *Escherichia coli*.

### Mycobacterial culturing conditions

All *M. abscessus* strains were maintained in Middlebrook 7H9 broth (271310, BD Diagnostics, Franklin Lakes, NJ, USA) with 0.2% (v/v) glycerol (GX0185, Supelco, Bellefonte, PA, USA), 0.05% (v/v) Tween-80 (P1754, MilliporeSigma, Burlington, MA, USA), and 10% (v/v) oleic acid-albumin-dextrose-catalase (OADC) (90000-614, VWR, Radnor, PA, USA). OADC and Tween-80 were omitted from medium used for experiments. *M. abscessus* cultures were shaken at 150 rpm at 37°C.

### Plasmid construction

Oligonucleotides used for all plasmid and strain construction are listed in Supplemental Table 3. *E. coli* cells were made competent using rubidium chloride and transformed according to standard protocols^105^.

pL5-UV15-TetO-bioA plasmid was produced by PCR amplification of *MAB_2688c* with 20 bp complementarity to pL5 PTetO Msm PonA1 truncation A-FLAG clone 1^106^ using Phusion High-Fidelity Polymerase (M0530, NEB, Ipswich, MA, USA), followed by isothermal assembly^107^ into NdeI (R0111, NEB) and HindIII-HF (R3104, NEB) digested pL5 PTetO Msm PonA1 truncation A-FLAG clone 1.

pMV306G13+Lux+zeo was constructed by replacing the L5 integrase, AttP site, and aminoglycoside-3’-phosphotransferase (*aph*) kanamycin resistance gene of pMV306G13+Lux^49^ with the Tweety integrase and AttP site as well as the zeocin resistance gene using isothermal assembly^107^.

### M. abscessus transformation

*M. abscessus* was grown to OD600 = 0.5 then washed 3 times at 22°C by pelleting 5000 x g for 7 minutes then resuspending in the initial culture volume of 10% glycerol. After the final wash, cells were resuspended in 1/100^th^ the initial culture volume of 10% glycerol. 50 µL of this final mixture was combined with 100 ng DNA in 1 µL water and incubated at 22°C for 5 minutes. This mixture was transferred to a 2 mm electroporation cuvette (89047-208, VWR), and electroporated at 2500 V, 125 Ω, 25 μF using an ECM 630 electroporator (45-0651, BTX, Holliston, MA, USA). 1 mL 7H9 broth was added to the electroporated cells, and cells were incubated shaking at 150 rpm for 4 hr at 37°C. 100 µL of this mixture was spread on 7H10 + 0.5% (v/v) glycerol + 10% (v/v) OADC agar plates using 4 mm borosilicate glass beads, and plates were incubated at 37°C for 4 days.

### Strain construction

*ΔbioA* pMV306G13+Lux+zeo strains were generated by recombineering^108^ knockout of *MAB_2688c* (*bioA*). *M. abscessus* ATCC19977 was transformed with 100 ng pNit-RecET^108^, a plasmid which contains a nitrile-inducible recombinase as well as the counter-selection SacBR genes^108^, and selected on 7H10 + 0.5% (v/v) glycerol + 10% (v/v) OADC agar plates containing 50 µg/mL kanamycin sulfate (K4000, MilliporeSigma). Successful transformants were identified by colony PCR using oligonucleotides MRS01 and MRS02. To generate the recombineering template, 3 fragments were produced with 20 bp overlaps by PCR amplification with Phusion High-Fidelity Polymerase using the indicated primer pairs: a fragment 500 bp upstream of MAB_2688c (MRS03 + MRS04) using *M. abscessus* genomic DNA as a template, the zeocin resistance cassette flanked by loxP sites using pKM-lox-zeo as a template^108^ (MRS05 + MRS06), and a fragment 500 bp downstream of MAB_2688c (MRS07 + MRS08) using *M. abscessus* genomic DNA as a template. Those three fragments were joined with a NotI-HF (R3189, NEB) and NdeI digested vector, pL5 PTetO Msm PonA1 truncation A-FLAG clone 1^106^, using isothermal assembly^107^. The linear recombineering product was PCR amplified from the resulting plasmid using Phusion High-Fidelity Polymerase and primers MRS09 + MRS10 and PCR purified using the Monarch PCR & DNA Cleanup Kit (T1030, NEB). *M. abscessus* pNit-RecET was grown to OD600 = 0.8, then 1 µM isovaleronitrile (308528, MilliporeSigma) was added to the culture for 8 hr, followed by addition of 200 mM glycine for 14 hr. *M. abscessus* pNit-RecET was then transformed with 1 µg linear recombineering fragment as described above, with the following modifications: final resuspension of electrocompetent cells was in 1/10^th^ the initial culture volume, electroporator settings were 2500 V;1000 Ω;25 μF, and cultures were allowed to recover at 37°C for 8 hr prior to plating. Successful recombineering deletions were confirmed by colony PCR (Supplemental Figure 2D) using primers MRS09 + MRS10. *bioA::zeoR* strains were passaged on 7H10 + 0.5% (v/v) glycerol + 10% (v/v) OADC agar plates containing 3% sucrose to select for bacteria that lost the episomal plasmid pNit-RecET and its SacBR gene. Successful loss of pNit-RecET was identified by failure to grow on 7H10 + 0.5% (v/v) glycerol + 10% (v/v) OADC agar plates containing 50 µg/mL kanamycin along with ability to grow on 7H10 + 0.5% (v/v) glycerol + 10% (v/v) OADC agar plates containing 100 µg/mL zeocin. *M. abscessus bioA::zeoR* strains were then transformed with pCreRec-SacBR-kan^108^ to excise the *zeoR* gene and produce *ΔbioA* strains, which were confirmed by colony PCR with primers MRS09 + MRS10 (Supplemental Figure 2D). *ΔbioA* strains were made luminescent by transformation with 100 ng pMV306G13+Lux+zeo and selection on 7H10 + 0.5% (v/v) glycerol + 10% (v/v) OADC agar plates containing 100 µg/mL zeocin (R25001, Thermo Fisher Scientific, Waltham, MA, USA). *bioA* rescue constructs were generated by transforming *ΔbioA* pMV306G13+Lux+zeo strains with pUV15-Tet-bioA and selecting on 7H10 + 0.5% (v/v) glycerol + 10% (v/v) OADC agar plates containing 50 µg/mL kanamycin sulfate and 100 µg/mL zeocin.

Luminescent strains were generated by transforming 100 ng pMV306G13+Lux^49^ into electrocompetent *M. abscessus* and selecting on 7H10 + 0.5% (v/v) glycerol + 10% (v/v) OADC agar plates containing 50 µg/mL kanamycin. Colonies were checked for luminescence using the chemiluminescence setting of a c300 Gel Imaging System (Azure Biosystems, Dublin, CA, USA).

Fluorescent *M. abscessus* was generated by transformation with 100 ng plasmid pL5-MOP-mScarlet containing mScarlet^50^ driven by a Mycobacterial Optimized Promoter (MOP)^109^, followed by selection on 7H10 + 0.5% (v/v) glycerol + 10% (v/v) OADC agar plates containing 50 µg/mL kanamycin.

### Colony PCR

100 µL saturated *M. abscessus* culture was pelleted 10,000 x g 1 min. The pellet was resuspended in 10 µL sterile water, then 1 µL was transferred to a PCR tube containing 500 nM of each forward and reverse primer and 1X GoTaq mix (M7123, Promega, Madison, WI, USA) in 20 µL total volume. PCR tubes were incubated at 95°C for 30 minutes to sterilize cultures, then PCR was performed according to manufacturer’s recommendations.

### Air-liquid interface culture

NuLi-1 immortalized bronchial epithelial cells^48^(ATCC CRL-4011) were expanded in bronchial epithelial growth medium (BEGM)^110^ (See Supplemental File 1 for formulation) on collagen coated 75 cm^2^ flasks (353136, Corning Inc., Corning, NY, USA). Cells were washed with phosphate buffered saline (PBS) (10010031, Thermo Fisher Scientific), detached by addition of 0.25% trypsin-0.02% EDTA (59428C, MilliporeSigma) followed by 10 minute incubation at 37°C, and counted using a Countess II FL cell counter (Thermo Fisher Scientific) after addition of trypan blue (T8154, MilliporeSigma) to assess viability. Cells were plated in either 24-well (62406-173, VWR) or 6-well (62406-171, VWR) transwell inserts coated with collagen at densities of 90,000 cells per 24-well insert or 800,000 cells per 6-well insert with BEGM medium added to both the basal and apical compartments. Cells were allowed to expand in BEGM for 3 days to ensure formation of a confluent monolayer, then BEGM was removed from both compartments and replaced with air-liquid interface culture (ALI) medium^110^ (See Supplemental File 1 for formulation) only in the basal compartment. 24-well cultures were provided with 0.8 mL basal medium, and 6-well cultures were provided 3 mL basal medium. ALI cultures were maintained for 14 days, changing basal media and aspirating apical liquid every 2 days. For all cell culture, cells were maintained in a humidified incubator with 5% CO_2_ at 37°C. NuLi-1 cells routinely tested negative for mycoplasma infection using MycoAlert Mycoplasma Detection Kit (LT07-418, Lonza Group AG, Basel, Switzerland) according to the manufacturer’s instructions.

### Collagen coating plates

Human placental collagen (C7521, MilliporeSigma) was resuspended at 0.1 mg/mL in PBS, left overnight at 4°C to allow collagen to dissolve, and filtered through a 0.22 µm filter (SE1M179M6, MilliporeSigma). Sterile filtering reduces collagen concentration to an unknown degree, so collagen was always filtered using same type of filter. The following volumes of collagen solution were added to plates or transwells: 3 mL for 75 cm^2^ flasks, 1 mL for 6-well transwell, and 0.3 mL for 24-well transwell. Collagen was left on plates and transwells overnight at 4°C, then aspirated from plates the following day.

### Lung infection model

Mature air-liquid interface cultures were infected with the indicated amounts of *M. abscessus* by adding *M. abscessus* resuspended in PBS to the apical compartment of the culture. *M. abscessus* was suspended in 50 µL PBS per well for 24-well cultures and 500 µL PBS per well for 6-well cultures. To ensure that the air-liquid interface was maintained, cultures were infected, incubated for 4 hours, then excess liquid was removed from the surface. This process consistently removed approximately 80% of the bacteria (Supplemental Figure 1D). MOI noted on all graphs represents the targeted MOI after removal of excess medium. Infected cultures were sealed with Breathe-Easy membranes (Z380059, MilliporeSigma) and incubated in a humidified incubator with 5% CO_2_ at 37°C. Luminescent *M. abscessus* measurements were taken using a Spark 10M plate reader (Tecan, Mannedorf, Switzerland) in the same culture plates used for infection. To avoid luminescence bleed-through between wells, cultures were spaced at intervals across the culture plate. Lung cell viability was monitored by lactate dehydrogenase release using the LDH-Glo Cytotoxicity assay (J2380, Promega) according to manufacturer’s instructions.

### Fluorescein permeability assay

Sodium fluorescein (46960, MilliporeSigma) was added to the apical compartment of air-liquid interface cultures, or control collagen-coated wells with no cells. Equal volumes of apical and basal liquid were transferred to a black 96-well plate (3915, Corning), and fluorescence was measured in a Tecan Spark 10M plate reader with an excitation wavelength of 482 nm and an emission wavelength of 527 nm. Monolayer permeability represents the ratio of fluorescein in the basal compartment compared to the apical compartment normalized to the ratio present in collagen-coated, empty wells.

### Fluorescence microscopy

Lung cells were fixed directly to transwells by treatment with 4% paraformaldehyde (15710, Electron Microscopy Sciences, Hatfield, PA) in PBS for 1 hr at 22°C. After fixation, cells were stored in PBS at 4°C until stained. Fixed cells attached to the transwell membrane were cut out of the transwell and transferred to a 1.5 mL microcentrifuge tube. Cells were permeabilized by treatment with 250 µL PBS + 1 mM CaCl_2_ (0556, VWR), 1 mM MgCl_2_ (M8266, MilliporeSigma), and 0.2% (v/v) Triton X-100 (T8787, MilliporeSigma) for 15 minutes at 22°C, then washed 3x with PBST (PBS + 1 mM MgCl_2_ + 1 mM CaCl_2_ + 0.1% Tween 20 (P1379, MilliporeSigma)). For f-actin staining, membranes were incubated on a rocker in 100 µL phalloidin-iFluor 488 (ab176753, Abcam, Cambridge, UK) diluted 1:1000 in PBS + 1% bovine serum albumin (A9647, MilliporeSigma) for 1 hr at 22°C. For MUC5AC staining, permeabilized membranes were incubated on a rocker 1 hr at 22°C in PBST + 1% bovine serum albumin + 10% goat serum (ab7481, Abcam), then incubated 16 hr at 4°C in anti-MUC5AC antibody (ab3649, Abcam) diluted 1:20 in PBST + 1% bovine serum albumin + 10% goat serum. Membranes were then washed 3x with PBST and incubated 1 hr at 22°C in 100 µL anti-mouse IgG AlexaFluor 594 secondary antibody (ab150116, Abcam) diluted 1:200 in PBST + 1% bovine serum albumin + 10% goat serum. After staining, all membranes were washed 3x in PBST. Where relevant, cells were incubated 1 minute at 22°C in 100 µL 1 µg/mL 4′,6-diamidino-2-phenylindole (DAPI) (D9542, MilliporeSigma) in PBS, then washed once in PBS. Transwell membranes were transferred to glass slides (16004430, VWR), 10 µL n-propyl gallate solution (50 mg/mL n-propyl gallate (MP210274780, Thermo Fisher Scientific) and 16.3 mg/mL Tris base (648310, MilliporeSigma) dissolved in 70:30 glycerol:PBS) was added to the slides as an anti-fade reagent, and slides were covered with 22 mm x 22 mm, 1.5 thickness glass cover slips (3406, Erie Scientific, Ramsey, MN, USA). For widefield images (Supplemental Figure 1B), slides were imaged on an inverted Nikon TI-E microscope at the indicated magnification. Phalloidin-iFluor 488 was excited at 470 nm, anti-mouse AlexaFluor 594 was excited at 555 nm, and DAPI was excited at 395 nm. For confocal images (Figure 1B, Supplemental Figure 1G-H), images were collected on a Zeiss LSM980 single point scanning confocal microscope with a 63x 1.4 NA oil-immersion objective and a 1024×1024 pixel frame size. Phalloidin-iFluor 488 was excited at 488 nm and emission was monitored over the range 482-677 nm. mScarlet-expressing *M. abscessus* was excited at 561 nm and emission was monitored over the range 569-700 nm. DAPI was excited at 405 nm and emission was monitored over the range 378-686 nm. For orthogonal view (Supplemental Figure 1H), z-stack images were taken at increments of 0.25 µm. Images were processed using Fiji^111^ running ImageJ v1.53q.

### Scanning electron microscopy

Lung cells were fixed directly to transwells for 18 hr at 22°C in a mixture of 1.25% formaldehyde, 2.5 % glutaraldehyde and 0.03% picric acid in 0.1 M Sodium cacodylate buffer, pH 7.4. Fixed cells were washed with 0.1 M sodium cacodylate buffer and post-fixed with 1% osmium tetroxide in 0.1 M sodium cacodylate buffer for 2 hours at 22°C. Cells were then rinsed in ddH_2_O and dehydrated through a series of ethanol (30%, 50%, 70%, 95%, (2x)100%) for 15 minutes per solution. Dehydrated cells were then placed in a 1:1 solution of hexamethyldisilazane (HMDS) and 100% ethanol for 1 hour at 22°C, then washed 2x 30 minutes at 22°C with 100% HMDS. Samples were left in a fume hood to air dry 18 hr at 22°C, then mounted on aluminum stages with carbon dots and coated with platinum (6 nm) using a Leica EM ACE600 Sputter Coater. The dried samples were observed in a Hitachi S-4700 Field Emission Scanning Electron Microscope (FE-SEM) at an accelerating voltage of 3kV.

### Luminescence growth curves

*M. abscessus* was thawed and grown to saturation in 7H9 broth with 0.2% (v/v) glycerol, 0.05% (v/v) Tween-80, and 10% (v/v) OADC, then diluted back and grown overnight to an OD600 = 0.5-0.8. Based on OD600 measurements, approximately 10,000 colony forming units were pelleted and resuspended in PBS and then plated in each well of a white, 96-well plate (655074, Greiner Bio-One, Frickenhausen, Germany) in 100 µL of relevant medium. For rescue experiments, growth medium was prepared by adding small volumes of concentrated stock solutions of the desired compound. Stock solutions were made as follows: 1 M stock solution of sodium acetate (S2889, MilliporeSigma) in water, 200 mM stock of sodium propionate (P1880, MilliporeSigma) in water, 50 mM stock solution of octadec-9-enoate (O7501, MilliporeSigma) in water, 50 mM stock solutions of hexadecanoic acid (P5585, MilliporeSigma), hexadec-9-enoic acid (P9417, MilliporeSigma), octadecanoic acid (S4751, MilliporeSigma), and pentadecanoic acid (P6125, MilliporeSigma) in ethanol (MACR6777-16, VWR), and 20% tyloxapol (T8761, MilliporeSigma) in water. Pure benzyl alcohol (24122, MilliporeSigma) was added directly to medium. For dialyzed medium experiments, basal medium from air-liquid interface cultures was taken after 48 hr incubation, then dialyzed against ALI medium using a Pur-A-Lyzer Maxi Dialysis Kit (PURX35015, MilliporeSigma) according to manufacturer’s instructions. For all growth curves, plates were sealed with Breathe-Easy membranes and incubated at 37°C without shaking.

### Transposon library production

To create a transposon mutant library of strain T35, 100 mL of T35 culture was grown to an OD600 of 1.5. Cells were washed twice with 50 mL MP Buffer (50mM Tris-HCl pH 7.5, 150 mM NaCl, 10 mM MgSO_4_, 2 mM CaCl_2_) and resuspended in 10 mL MP Buffer. 2 x 10^11^ phage forming units of temperature sensitive φMycoMarT7 phage^112^ carrying the Himar1 transposon^113^ were added to bacteria. Phage and bacterial cultures were incubated at 37°C for 4 hr with shaking. Transduced cultures were pelleted at 3200 x g for 10 min at 22°C and then resuspended in 12 mL of PBS + 0.05% Tween 80. Cultures were titered by plating on 7H10 + 0.5% (v/v) glycerol + 10% (v/v) OADC agar plates supplemented with 100 µg/mL kanamycin sulfate. 150,000 bacterial mutants were plated onto 7H10 + 0.5% (v/v) glycerol + 10% (v/v) OADC + 0.1% (v/v) Tween 80 + 100 µg/mL kanamycin sulfate agar plates and grown for 4 days at 37°C. The resulting mutant library was harvested and stored in 7H9 + 10% glycerol at −80°C.

### Transposon library selection in lung infection model

Transposon mutant libraries were inoculated at a final MOI = 1 onto mature air-liquid interface cultures of NuLi-1 cells grown in 6-well transwells as described above. Libraries were also inoculated into 3 mL ALI medium (Supplemental File 1) in 6-well plates without transwells for the “Tissue culture medium” condition. 3 biological replicates of the lung infection and 3 biological replicates of the tissue culture medium condition were inoculated. After 48 hr, M. abscessus was harvested from lung cultures by adding 500 µL PBS to the apical surface, scraping the apical surface of the transwell with a scraper (734-2602, VWR), and transferring PBS to a microcentrifuge tube. Bacteria were pelleted 5000 x g for 5 minutes at 22°C, resuspended in 7H9 + 0.2% (v/v) glycerol + 0.05% (v/v) Tween-80 + 10% (v/v) OADC, mixed 1:1 with 50% glycerol, and frozen at −80°C until titering. Cultures were titered by plating on 7H10 + 0.5% (v/v) glycerol + 10% (v/v) OADC agar plates supplemented with 100 µg/mL kanamycin sulfate. Approximately 150,000 bacterial mutants for each condition were plated across 6 245 mm x 245 mm (431111, Corning) 7H10 + 0.5% (v/v) glycerol + 10% (v/v) OADC + 0.1% Tween 80 + 100 µg/mL kanamycin sulfate agar plates and grown for 4 days at 37°C. All 6 plates for each biological replicate were combined by scraping into a 50 mL conical tube containing 5 mL 7H9 + 0.2% (v/v) glycerol + 0.05% (v/v) Tween-80 + 10% (v/v) OADC and 5 mL 50% glycerol. Post-selection libraries were then frozen in 2 mL aliquots at −80°C until gDNA isolation.

### Genomic DNA Extraction

gDNA was isolated as previously described^25^. 2 mL post-selection transposon mutant libraries were pelleted, resuspended in TE Buffer (10 mM Tris HCl pH 7.4, 1 mM EDTA pH 8), and transferred to 2 mL tubes containing 0.1 mm silica beads (116911500, MP Biomedicals, Irvine, CA) as well as 600 µL 25:24:1 phenol:chloroform:isoamyl alcohol (P3803, MilliporeSigma). Samples were homogenized using Bead Bug 3 Microtube Homogenizer (D1030, Benchmark Scientific, Sayreville, NJ, USA) 4 x 45 s at 4000 rpm. Samples were cooled on ice for 45 s between each successive round of homogenization. Samples were pelleted 21,130 x g 10 minutes at 22°C, then aqueous layer of supernatant was combined with 1 volume 25:24:1 phenol:chloroform:isoamyl alcohol and incubated on a rocker 1 hr at 22°C. Samples were then transferred to pre-pelleted MaXtract High Density phase-lock tubes (129065, Qiagen, Hilden, Germany), centrifuged 1500 x g 5 minutes at 4°C, re-extracted by adding ½ volume chloroform (193814, MP Biomedicals), and centrifuged 1500 x g 5 minutes at 4°C. Upper aqueous layer was transferred to a new MaXtract High Density phase-lock tube and samples were incubated shaking at 150 rpm 1 hr at 22°C with RNase A (EN0531, Thermo Fisher Scientific) added to a final concentration of 25 µg/mL. Samples were then re-extracted with 1 volume 25:24:1 phenol:chloroform:isoamyl alcohol, centrifuged 1500 x g 5 minutes at 4°C, extracted with ½ volume chloroform, and centrifuged 1500 x g 5 minutes at 4°C. The aqueous phase was transferred to a conical tube, and DNA was precipitated by adding 1/10^th^ volume 3 M sodium acetate pH 5.2 and 1 volume isopropanol (3032-16, VWR) then incubating at 22°C for 18 hr. Pellet was washed 3x with 70% ethanol, dried 10 minutes to remove residual ethanol, then resuspended in 1 mL nuclease free water.

### Transposon sequencing, mapping, and analysis

Chromosomal-transposon junctions were amplified following established protocols^114^. These amplicons were sequenced using an Illumina NextSeq 500 sequencer, and reads were mapped to the T35 genome using TRANSIT Pre-Processor and analyzed using TRANSIT^115^. Insertion counts were normalized to trimmed total reads, and comparisons of gene essentiality between conditions were performed with permutation-based resampling analysis^115^. Multiple comparison-adjusted p-values were determined using the Benjamini-Hochberg method.

### Biotin quantitation

Culture medium was sampled from the lung infection model, the liquid was centrifuged at 5000 x g for 5 minutes to pellet any cells, and the supernatant was transferred to a new microcentrifuge tube and incubated at 85°C for 1 hr to ensure any remaining bacteria were heat-killed. Standards were also incubated at 85°C for 1 hr. Biotin was quantitated using a competitive enzyme-linked immunosorbent assay (ELISA) kit (K8141, Immundiagnostik, Bensheim, Germany) according to manufacturer’s instructions. Samples were diluted 1:75 in the ELISA kit sample dilution buffer. Absorbance was measured at 450 nm using a Tecan Spark 10M plate reader.

### Protein isolation and western blot

Protein was isolated from *M. abscessus* by pelleting bacteria 3200 x g for 10 minutes at 4°C, resuspending in Tris buffered saline (TBS) (28358 Thermo Fisher Scientific) + protease inhibitor (11873580001, MilliporeSigma) (0.5 tablet per 10 mL TBS), transferring to 2 mL tubes with 0.1 mm silica beads (116911500, MP Biomedicals), and homogenizing using a Bead Bug 3 Microtube Homogenizer 4 x 45 seconds at 4000 rpm with 2 minutes of incubation on ice between rounds of homogenization. Homogenized samples were pelleted 21,130 x g for 5 minutes at 4°C, and the supernatant was heat-killed by incubation at 80°C for 20 minutes. Protein abundance was quantitated by absorbance at 280 nm using a Nanodrop 1000 spectrophotometer (Thermo Fisher Scientific), and samples were normalized to 2.5 mg/mL by dilution in TBS. After normalization, remaining DNA was digested by addition of TURBO DNase buffer (AM2238, Thermo Fisher Scientific) (final concentration of 10%) and TURBO DNase (final concentration of 2%) followed by incubation at 37°C for 15 minutes. Samples were mixed with 4X LDS NuPage sample buffer to a final concentration of 1X and dithiothreitol (71003-396, VWR) to a final concentration of 50 mM. Samples were incubated at 70°C for 10 minutes, then 7 µg of protein along with PageRuler Prestained ladder 10 kDa to 180 kDa (26616, Thermo Fisher Scientific) was loaded on a NuPage 4-12% gradient Bis-Tris pre-cast SDS-PAGE gel (NP0321, Thermo Fisher Scientific), which was electrophoresed at 115 V for 90 minutes. Proteins were transferred to a PVDF membrane (1704156, Bio-Rad Laboratories, Hercules, CA) using TransBlot Turbo Transfer System (Bio-Rad) on the Mixed MW setting. Membranes were blocked by incubating in TBS + 0.1% Tween 20 (TBST) + 5% bovine serum albumin 1 hr at 22°C, and then were incubated with streptavidin-HRP (3999S, Cell Signaling Technology, Danvers, MA, USA) diluted 1:400,000 in TBST + 5% bovine serum albumin for 18 hr at 4°C. Membranes were washed 3x in TBST to remove unbound streptavidin-HRP and were developed by 1 minute incubation in 1 mL Azure Radiance Plus luminol/enhancer solution and 1 mL Azure Radiance Plus Peroxide Chemiluminescent Detection Reagent (AC2103, Azure Biosystems). Excess reagent was allowed to drain off of the membrane, and membranes were imaged using the chemiluminescence detector of a c300 Gel Imaging System (Azure Biosystems). After blotting, total protein was detected by staining membranes with SYPRO Ruby Protein Blot Stain (S11791, Thermo Fisher Scientific) according to manufacturer’s instructions. SYPRO Ruby staining was imaged using the Epi Blue setting of the c300 Gel Imaging System.

### Fatty acid methyl ester production

To avoid detergent contamination for mass spectrometry, *M. abscessus* ATCC19977 Tween-free glycerol stocks were generated by growing *M. abscessus* to saturation in 7H9 + 0.2% (v/v) glycerol + 10% (v/v) OADC, then freezing at −80°C in 25% glycerol in 7H9. From these stocks, luminescent *M. abscessus* was cultured in experimental medium to a luminescence value equivalent to OD600 = 0.6, then collected by centrifugation at 3200 x g for 7 minutes at 4°C. Cells were washed 2x in HPLC grade water (270733, MilliporeSigma), then resuspended in 1 mL HPLC grade water and transferred to a glass tube with PTFE-lined cap. Total lipids were isolated by Folch extraction^116^ by adding 24 mL 2:1:0.6 HPLC chloroform (C297-4, Thermo Fisher Scientific): HPLC methanol (A454-4, Thermo Fisher Scientific): HPLC water with 10 µg/mL butylated hydroxytoluene (B1378, MilliporeSigma) as an antioxidant and by then shaking the samples 18 hr at 22°C. Samples were centrifuged 1600 x g for 10 minutes at 22°C to separate layers, then the lower organic layer was transferred to a pre-weighed glass tube. Samples were dried under continuous flow of atmospheric air and were then weighed to determine yield. Fatty acid methyl esters were generated by acid-catalyzed methyl esterification. Samples were dissolved at 10 mg/mL in toluene, then 50 µL of resuspended sample was evaporated to dryness under continuous flow of nitrogen, and 450 µL 0.4 M HCl in methanol was added to the samples and incubated for 18 hr at 50°C. 250 µL of 5% NaCl in water and 250 µL hexanes were added, then samples were vortexed and set at 22°C to allow layers to separate. The upper hexane layer was transferred to glass insert GC/MS vials.

### Gas chromatography/mass spectrometry

1 µL of sample or of a fatty acid methyl ester standard (CRM18918, MilliporeSigma) was injected into a 30 m x 250 μm x 0.25 μm DB-FastFAME column (G3903-63011, Agilent Technologies, Santa Clara, CA, USA) using helium as a carrier gas at a constant pressure of 14 PSI. The GC oven temperature was held at 50°C for 30 s, then increased at a rate of 25°C/min to 194°C and held for 1 min. Temperature was then increased at a rate of 5°C/min to 245°C and held for 3 min. The mass spectrometer (MS) was operated using electron impact ionization at 70 eV, with the MS source held at 230°C and the MS quadrupole held at 150°C. Ions were detected in normal scanning mode over an m/z range of 104-412.

### GC/MS peak identification and quantitation

GC/MS peaks were identified using AMDIS^117^ by comparison to a fatty acid methyl ester standard (CRM18918, MilliporeSigma) or by predicted retention time based on equivalent chain length^118–120^ and by mass/charge ratio and fragmentation pattern. Double bond location could not be determined confidently based on the small degree of retention time separation for different positions, so unsaturated fatty acids are listed without assigning a position for double bonds. Methyl-group position for branched fatty acids could be determined for some species, and species that could represent multiple branched fatty acids are indicated as such in figure panels. Peaks identified may also contain chemically converted fatty acids produced by the process of acid-catalyzed methyl esterification. Peak areas were quantitated using El-MAVEN^121^, and samples were normalized by subtracting blank measurements and normalizing to total ion counts within each sample. For heat maps and principal component analysis, the mean normalized peak intensity for each metabolite was subtracted from the normalized peak intensity of each sample, then that value was divided by the standard deviation of the peak intensities for that metabolite across all samples. Plots were produced using Metaboanalyst 5.0^122^. Heat maps were produced using Euclidean distance measurement and Ward clustering. Samples were clustered for all heat maps, and metabolites were clustered for Supplemental Figure 4C-D.

### HPLC/MS lipidomics

*M. abscessus* was cultured from detergent-free stocks in biological quadruplicate, harvested, and washed as described for fatty acid methyl ester production. Total lipids were extracted by resuspending washed cell pellets in 1 mL HPLC grade methanol (A454-4, Thermo Fisher Scientific), transferring to a glass vial with PFTE cap, adding 3 mL HPLC grade methanol and 2 mL HPLC grade chloroform (C297-4, Thermo Fisher Scientific), then shaking at 22°C for 1 hr. Samples were centrifuged 750 x g for 30 minutes at 22°C, and supernatant was collected. Insoluble pellets were re-extracted with 6 mL 1:2 methanol:chloroform using the same method, and supernatants were pooled with those collected in the first extraction. Pooled supernatants were evaporated to dryness under continuous nitrogen flow. HPLC/MS was carried out using an Agilent 1260 Infinity LC system with a 6546 QTOF mass spectrometer using a previously published method^123^ with minor modifications. 10 μL of pooled dried lipids dissolved to 1 mg/mL in 70:30 (v/v) hexanes:isopropanol was injected into a normal-phase Inerstil Diol column (GL Sciences, Tokyo, Japan) and eluted with a binary gradient solvent system using 70:30 (v/v) hexanes:isopropanol as starting solvent and 70:30 (v/v) isopropanol:methanol as the final solvent. Both solvents had 0.1% formic acid and 0.05% aqueous ammonia added to improve ionization.

Extracted ion chromatograms from MassHunter software (Agilent Technologies) were generated for lipidomic analysis using the R package xcms^124^ for peak identification and alignment, statistical analysis using the linear model and Bayesian shrinkage of variance methods in the R package limma^125^, and data visualizations using base R. Code for R analyses is available by request. Visualization of mass spectra was carried out using MassHunter.

### Membrane fluidity measurements

*M. abscessus* cultures were grown in the indicated media to OD600 = 0.6, then laurdan (D250, Thermo Fisher Scientific) dissolved in dimethylformamide (DMF) was added to a final laurdan concentration of 10 µM and a final DMF concentration of 1% (v/v). Laurdan cultures were incubated 2 hr at 37°C with shaking and then collected by centrifugation at 3200 x g for 7 minutes at 22°C. Samples were washed 4x in the appropriate culture medium supplemented with 1% (v/v) DMF, then resuspended in 1/50 initial culture volume of appropriate culture medium + 1% (v/v) DMF. Samples were transferred to black 96-well plates (3915, Corning), and fluorescence was measured in a Tecan Spark 10M plate reader first at 23°C, then at 37°C after rapidly increasing the internal temperature of the plate reader. Laurdan was excited at 350 nm, and emission was monitored over a range from 440 nm to 490 nm. Fluorescence intensity measurements were converted into the laurdan generalized polarization (GP) metric^70^:

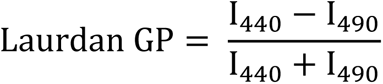

Higher values of laurdan GP indicate more ordered, less fluid membranes^67,70^.

### pH measurements

pH of medium was measured either using a potentiometric pH meter (30019028, Mettler Toledo, Columbus, OH, USA) or by measuring the ratio of 560 nm / 430 nm phenol red absorbance^100^ compared to a standard curve that was generated using a potentiometric pH meter.

---

**Supplemental Figure 1.**
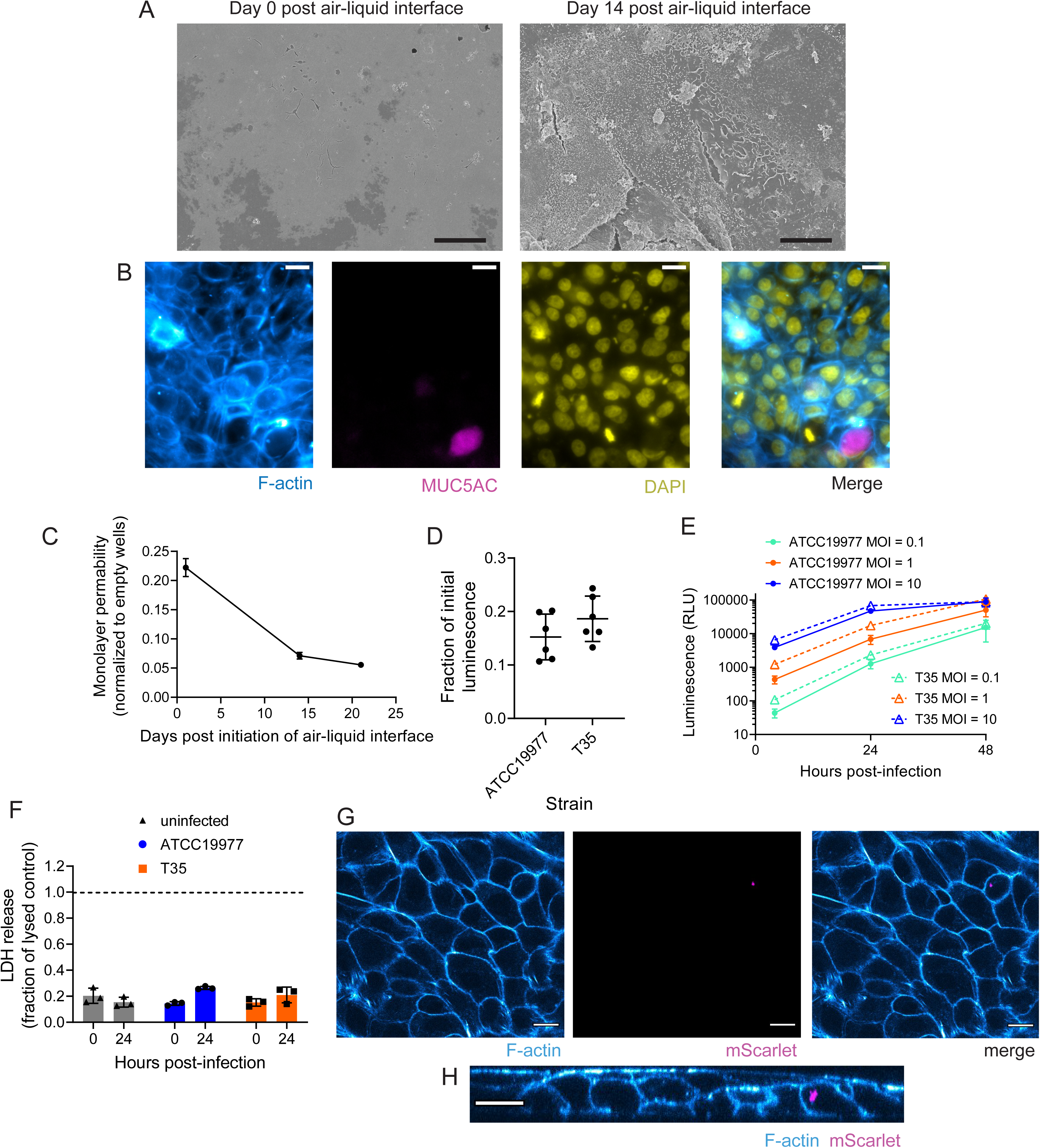
Air-liquid interface culture model system. **(A)** Scanning electron microscope images of apical surface of lung epithelial cells at 0 and 14 days after initiation of air-liquid interface. Images were obtained at 2000X magnification. Scale bar = 10 µm. **(B)** Widefield microscope images of NuLi-1 lung epithelial cells stained for F-actin, MUC5AC, and with DAPI to highlight nuclei. Images were obtained at 40X magnification. Scale bar = 20 µm. **(C)** Monolayer permeability as measured by amount of sodium fluorescein that penetrated through the epithelial layer at successive days after initiation of air-liquid interface. Data are normalized to empty, collagen-coated transwells and are presented as mean +/− SD. n = 3 biological replicates. **(D)** Fraction of *M. abscessus* remaining after aspiration of excess liquid to re-generate air-liquid interface for the *M. abscessus* type strain ATCC19977 and clinical isolate T35. Data are presented as individual values along with mean +/− SD. n=6 biological replicates. **(E)** Luminescence emitted by *M. abscessus* ATCC1997 or clinical isolate T35 in lung infection model infected at the indicated multiplicity of infection (MOI) over 48 hr of infection. Data are presented as mean +/− SD. n = 3 biological replicates per condition. **(F)** Lactate dehydrogenase (LDH) release from lung epithelial cells at 0 and 24 hr post-infection. LDH release is normalized to uninfected control cells that were lysed to release maximal LDH. Data are presented as individual values along with mean +/− SD. n = 3 biological replicates per condition. **(G)** Confocal microscope images of NuLi-1 lung epithelial cells infected with *M. abscessus* expressing mScarlet fluorescent protein, then washed to remove *M. abscessus* not internalized by lung cells and stained for F-actin. Images were obtained at 63X magnification. Scale bar = 15 µm. **(H)** Orthogonal view of cells pictured in **(G)**. Images were obtained at 63X magnification. Scale bar = 15 µm.

**Supplemental Figure 2.**
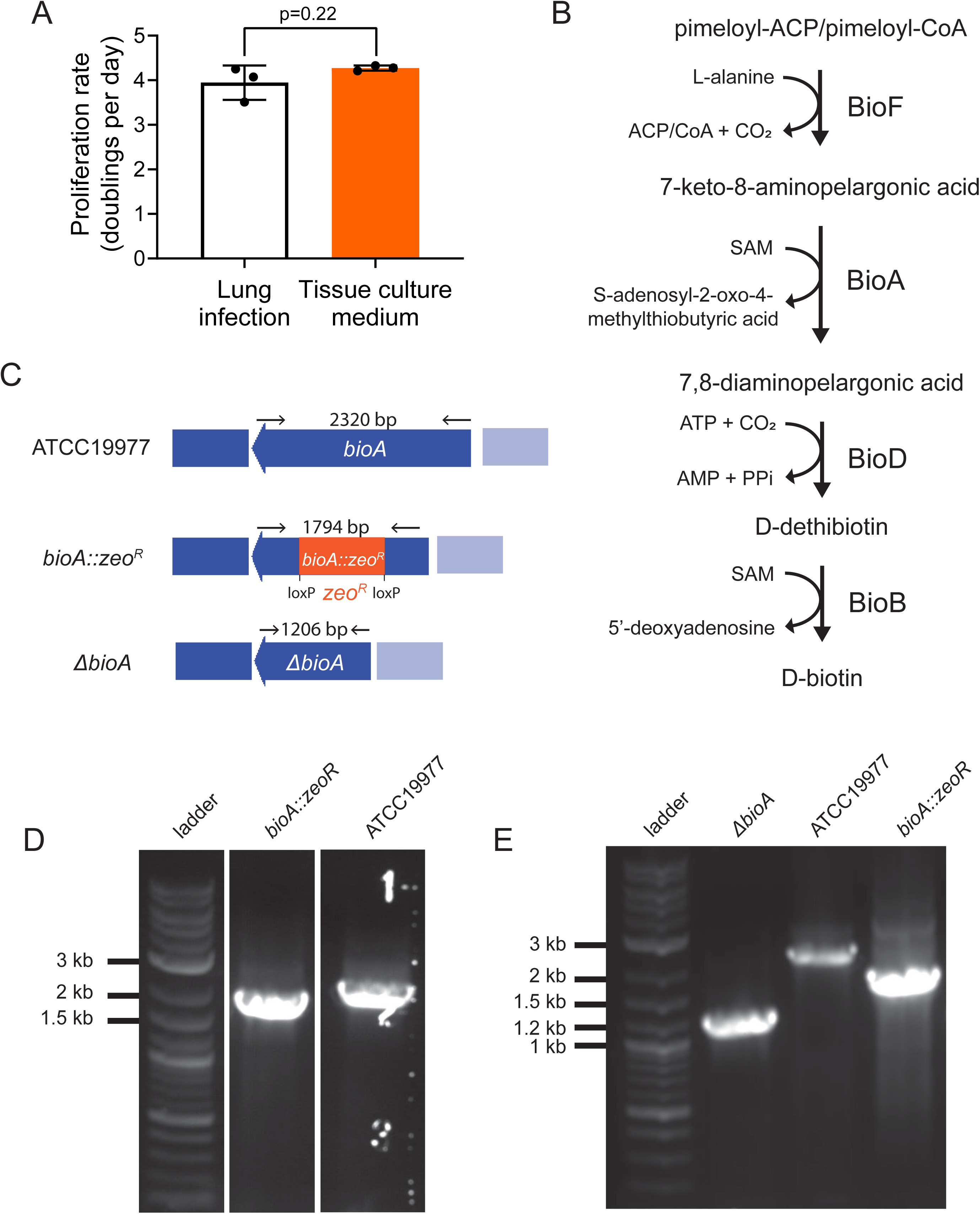
Development of biotin synthesis pathway knockouts. **(A)** Proliferation rate of *M. abscessus* ATCC19977 grown either in tissue culture medium or on the apical surface of air-liquid interface lung cultures. Data are presented as individual values along with mean +/− SD. n=3 biological replicates. p-value derived from unpaired, two-tailed t-test. **(B)** Schematic of biotin biosynthesis pathway. ACP: acyl carrier protein. CoA: coenzyme A. SAM: S-adenosyl methionine. ATP: adenosine triphosphate. AMP: adenosine monophosphate. PPi: inorganic phosphate. **(C)** Schematic of recombineering knockouts of *bioA*. zeoR: zeocin resistance cassette **(D)** Agarose gel electrophoresis of PCR products demonstrating insertion of zeoR into *bioA*. Expected PCR product sizes are indicated in **(C)**. **(E)** Agarose gel electrophoresis of PCR products demonstrating excision of zeoR from *bioA::zeoR*. Expected PCR product sizes are indicated in **(C)**.

**Supplemental Figure 3.**
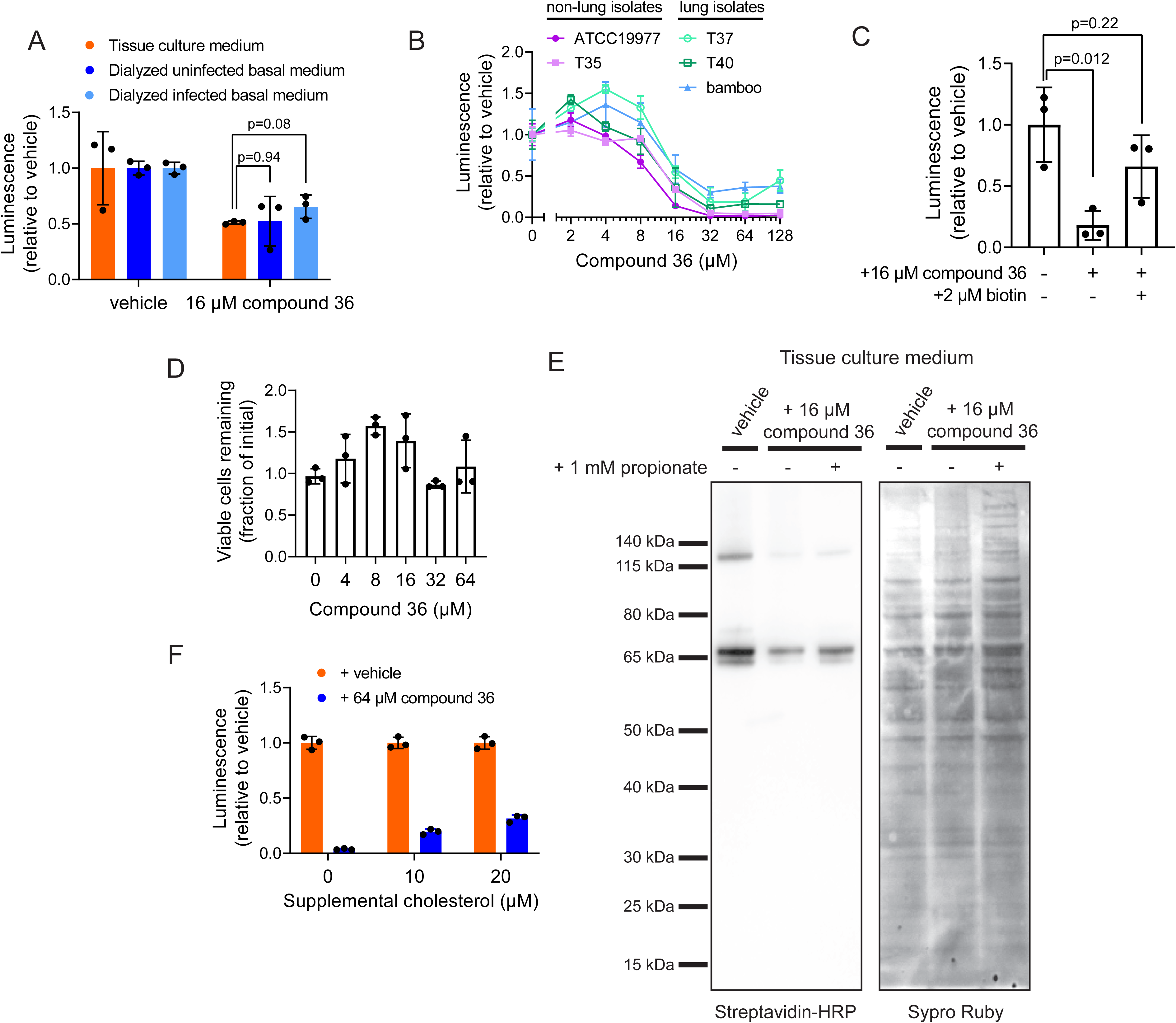
Characterization of BioA inhibitor compound 36 in *M. abscessus*. **(A)** Luminescence of *M. abscessus* ATCC19977 grown 48 hr with either vehicle or 16 µM compound 36 treatment. Media were either tissue culture medium or basal medium sampled from infected or mock infected air-liquid interface lung cultures dialyzed against tissue culture medium to replenish small molecules while retaining protein factors. Values are normalized within each condition to vehicle-treated. **(B)** Luminescence of the indicated *M. abscessus* clinical isolates grown in tissue culture medium for 48 hr with the specified final concentrations of the BioA inhibitor compound 36 in the medium. Values are normalized within each medium to the vehicle treated condition. **(C)** Luminescence of *M. abscessus* ATCC19977 grown in the lung infection model for 48 hr in the presence or absence of 16 µM compound 36 and/or 2 µM biotin added to the basal medium. Values are normalized to the vehicle treated condition. **(D)** Trypan blue measurement of viability of lung epithelial cells after 48 hr treatment with the indicated concentrations of compound 36. Values are normalized to vehicle treated condition. **(E)** Uncropped western blot (corresponding to Figure 3D) for total biotinylated protein in *M. abscessus* ATCC19977 grown in tissue culture medium with either vehicle or 16 µM compound 36 along with the indicated supplementation of propionate. SYPRO Ruby panel depicts total protein. **(F)** Luminescence of *M. abscessus* ATCC19977 grown in tissue culture medium for 48 hr with either vehicle or 64 µM compound 36 along with the indicated supplementation of cholesterol. Values are normalized within each condition to vehicle-treated. For all graphs, data are presented as mean +/− SD. n = 3 biological replicates. All p-values derived from unpaired, two-tailed t-tests.

**Supplemental Figure 4.**
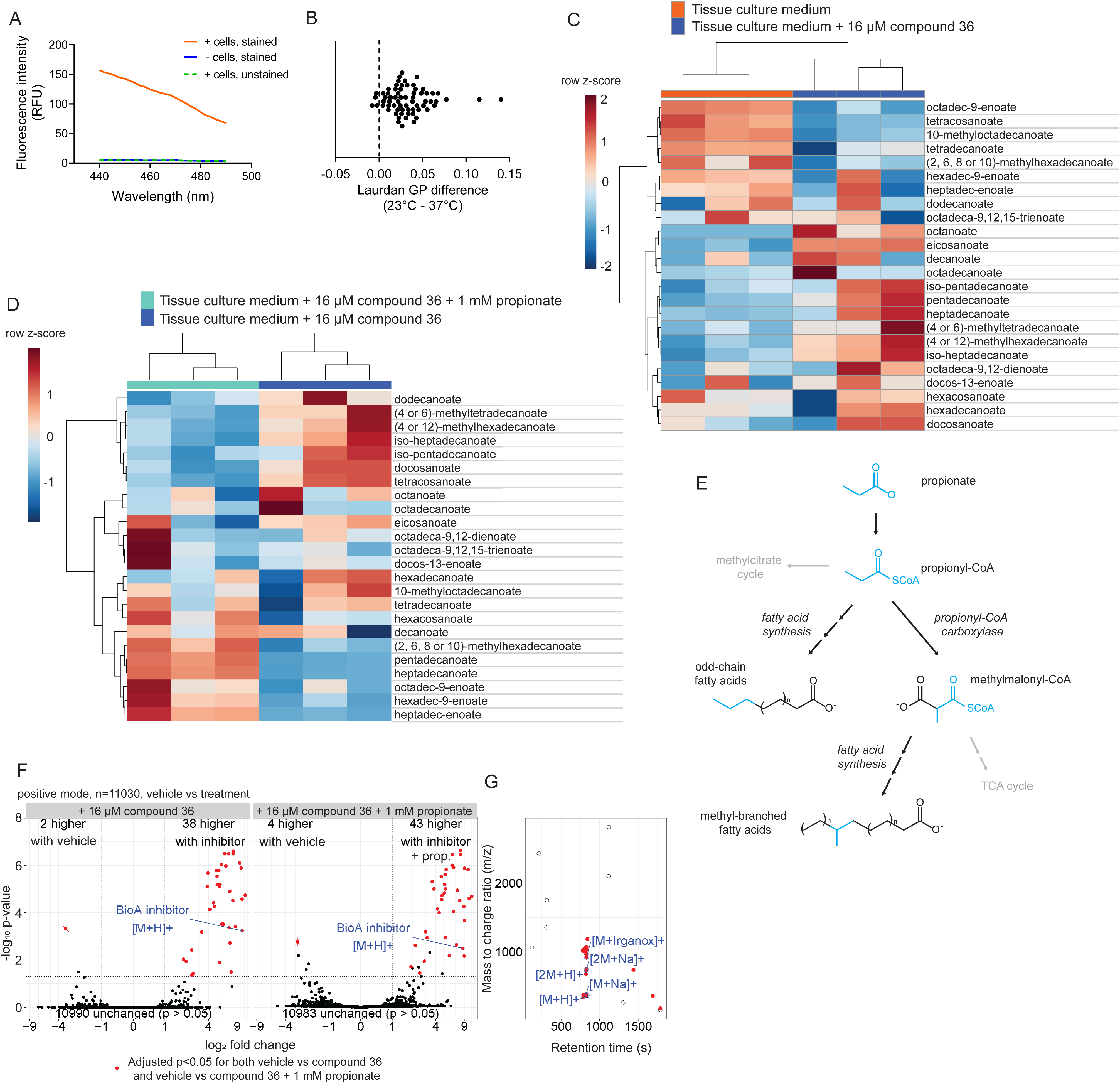
Altered biotin metabolism induces *M. abscessus* envelope remodeling **(A)** Fluorescence intensity scan from 440 nm to 490 nm from either laurdan stained (+cells, stained) or unstained (+cells, unstained) *M. abscessus* ATCC19977 samples or samples containing laurdan but no cells (-cells, stained). One representative sample is depicted for each condition. **(B)** Difference in laurdan generalized polarization (GP) between the same sample measured at 23°C, then rapidly shifted to 37°C and re-measured. n=65 biological replicates. **(C)** Heatmap depicting relative abundance of 24 fatty acid species measured by GC/MS in *M. abscessus* ATCC19977 grown 48 hr in tissue culture medium treated either with vehicle or 16 µM compound 36. Samples and fatty acid species are both hierarchically clustered. n=3 biological replicates. **(D)** Heatmap depicting relative abundance of 24 fatty acid species measured by GC/MS in *M. abscessus* ATCC19977 grown 48 hr in tissue culture medium treated with 16 µM compound 36 along with either vehicle or 1 mM sodium propionate. Samples and fatty acid species are both hierarchically clustered. n=3 biological replicates. **(E)** Schematic of propionate utilization. CoA: coenzyme A. TCA: tricarboxylic acid. **(F)** Volcano plots depicting log_2_-fold change in abundance versus significance for ‘molecular events’ with linked retention time, mass, and intensity representing potential lipid species detected by HPLC/MS. Molecular events detected in *M. abscessus* ATCC19977 grown 48 hr in tissue culture medium containing vehicle are contrasted against those detected in cells treated with 16 μM compound 36 (left) or with 16 μM compound 36 and 1 mM propionate (right). Peaks significantly changed (p < 0.05 after adjustment by the Benjamini-Hochberg method) in both contrasts (red circles) and a peak with the mass of compound 36 (blue outline) are indicated. Peak that is significantly depleted upon compound 36 treatment is depicted as an asterisk in both volcano plots. **(G)** Plot of retention time versus mass to charge ratio for all significantly changed peaks depicted in **(F)**, which clusters peaks by shared chemical properties. Peaks significant in both contrasts (red), a peak with the mass of compound 36 (blue, [M+H]+) and select alternate compound 36 adducts (blue) are indicated.

**Supplemental Figure 5.**
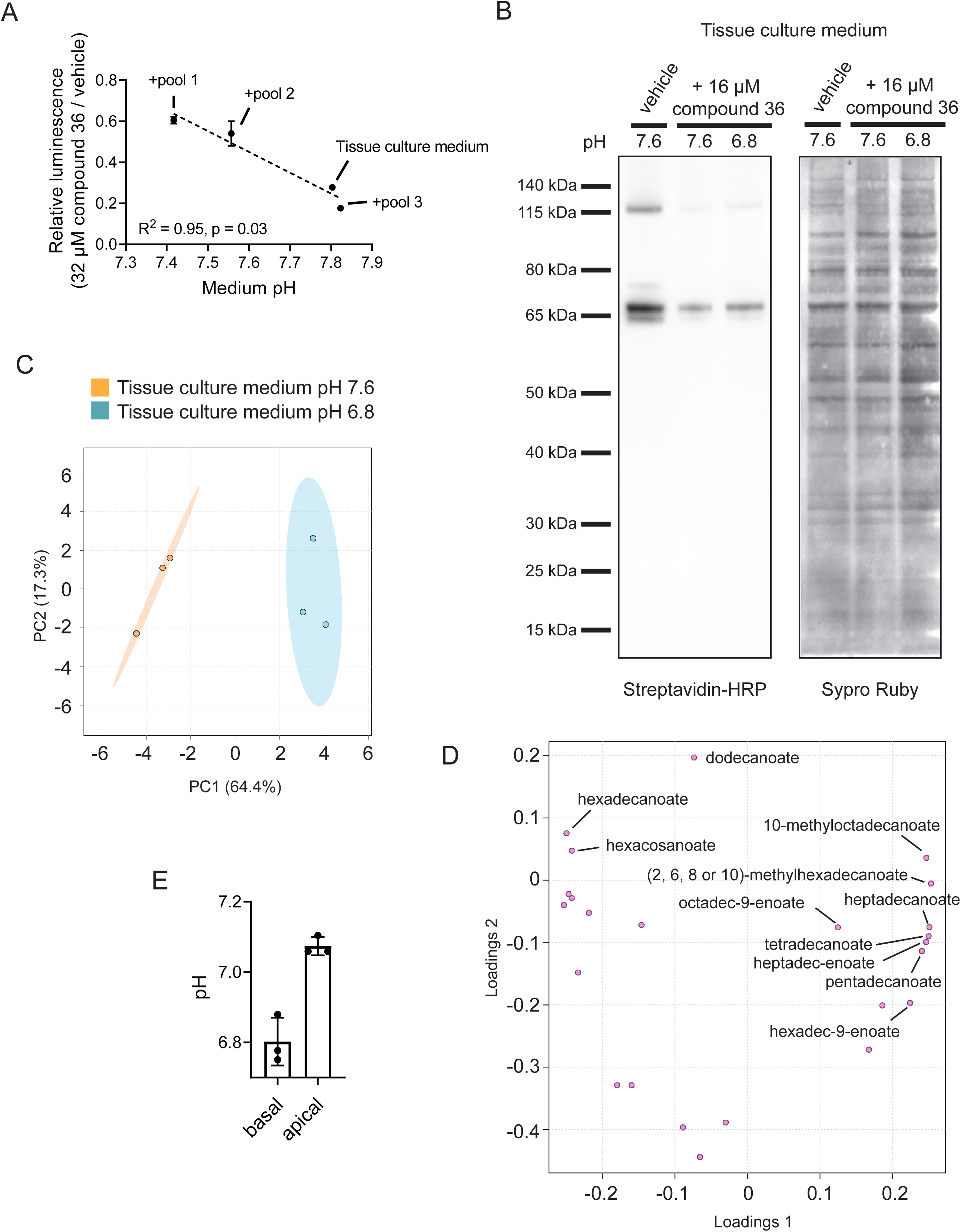
pH is a determinant of biotin demand and fatty acid composition. **(A)** Correlation between medium pH and sensitivity to biotin synthesis inhibition as measured by ratio of luminescence of *M. abscessus* ATCC19977 in tissue culture medium treated with 32 µM compound 36 compared to vehicle-treated after 48 hr. Medium pH changes are a secondary effect of adding pools of metabolites from mycobacterial medium to tissue culture medium (see Materials and Methods for composition of pools), and pH was measured by potentiometric pH meter. Data are presented as mean +/− SD. R^2^ and p-value derived from Pearson correlation. Line of best fit derived from simple linear regression. **(B)** Uncropped western blot (corresponding to Figure 5B) for total biotinylated protein in *M. abscessus* ATCC19977 grown in tissue culture medium adjusted to the indicated pH and treated with either vehicle or 16 µM compound 36. SYPRO Ruby panel depicts total protein. **(C)** Principal component analysis of *M. abscessus* ATCC19977 grown in tissue culture medium adjusted to the indicated pH based on GC/MS measurement of 24 fatty acid species. **(D)** Loading plot depicting individual fatty acid contributions to the principal components displayed in **(C)**. **(E)** pH of liquid sampled from the basal and apical surfaces of infected air-liquid interface lung cultures treated with 128 µM compound 36 as measured by phenol red absorbance. Data are presented as individual values along with mean +/− SD. n=3 biological replicates.

**Supplemental Table 1.** Library statistics for TnSeq.

| Library | Saturation (%) | Non-zero mean reads | Max read count | Skew |
| --- | --- | --- | --- | --- |
| T35 input | 55.6 | 34.0 | 1793 | 6.1 |
| T35 lung infection - 1 | 52.4 | 125.6 | 4327 | 3.6 |
| T35 lung infection - 2 | 38.1 | 94.7 | 2733 | 3.7 |
| T35 lung infection - 3 | 38.8 | 101.2 | 3246 | 3.9 |
| T35 tissue culture medium - 1 | 42.6 | 81.8 | 2513 | 4.0 |
| T35 tissue culture medium - 2 | 43.3 | 73.6 | 2489 | 4.4 |
| T35 tissue culture medium - 3 | 37.7 | 46.6 | 1599 | 4.0 |

**Supplemental Table 2.** Relative requirement for genes responsible for recycling propionate.

| Gene | Description | Insertion count log<sub>2</sub>-fold<br>change (lung infection<br>model / input library) | Adjusted p-value |
| --- | --- | --- | --- |
| *MAB_1462c* | Methylmalonyl-CoA<br>epimerase | -0.01 | 1.00 |
| *MAB_2711c* | Methylmalonyl-CoA<br>mutase subunit B | -0.35 | 1.00 |
| *MAB_2712c* | Methylmalonyl-CoA<br>mutase subunit A | -0.81 | 0.63 |
| *MAB_4616c* | Propionate regulator<br>(PrpR) | -0.37 | 1.00 |
| *MAB_4617* | Methylcitrate<br>dehydratase (PrpD) | -0.12 | 1.00 |
| *MAB_4618* | Methylcitrate lyase<br>(PrpB) | -0.52 | 1.00 |
| *MAB_4619* | Methylcitrate synthase<br>(PrpC) | 0.55 | 0.78 |

**Supplemental Table 3.**
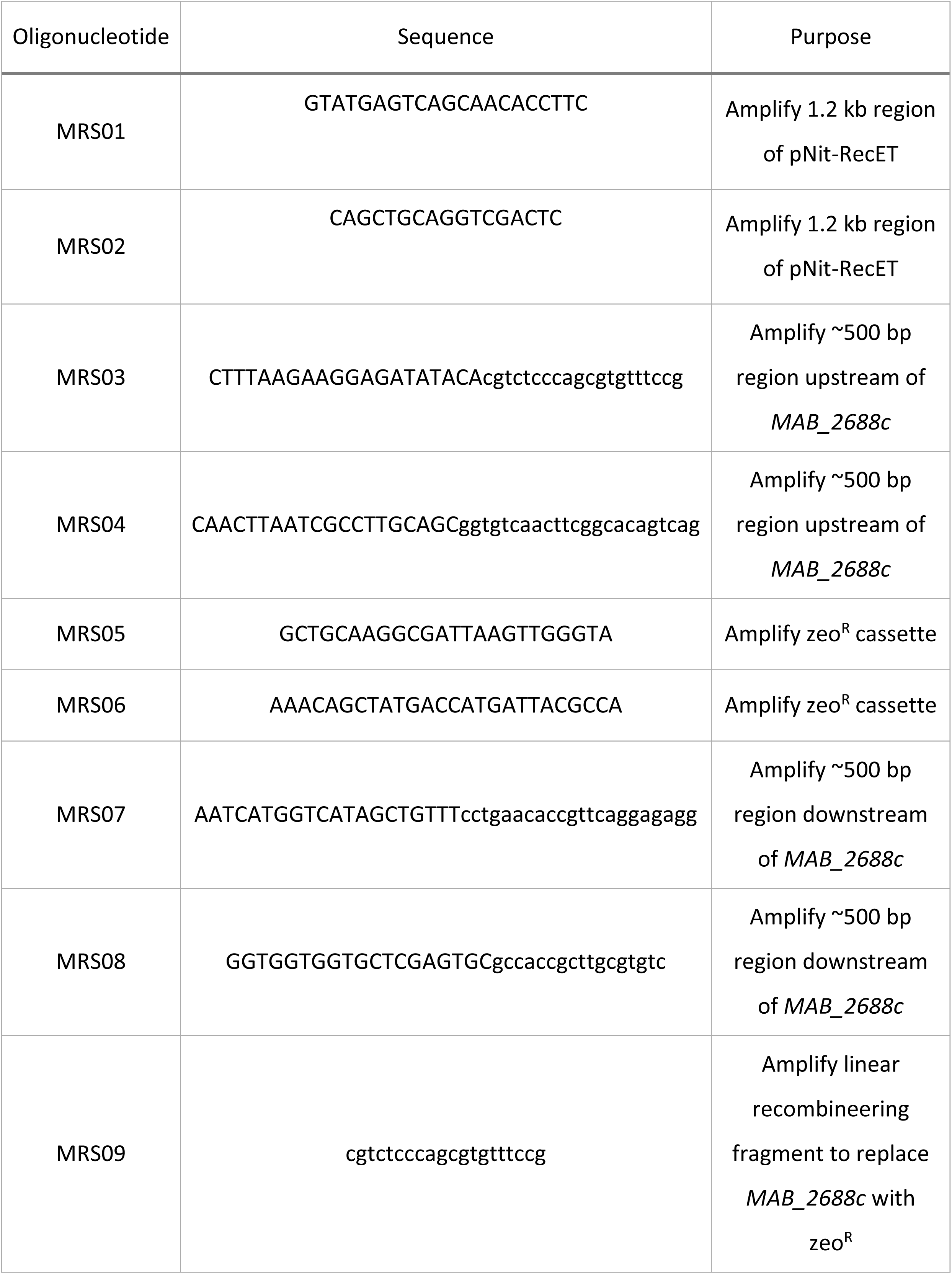
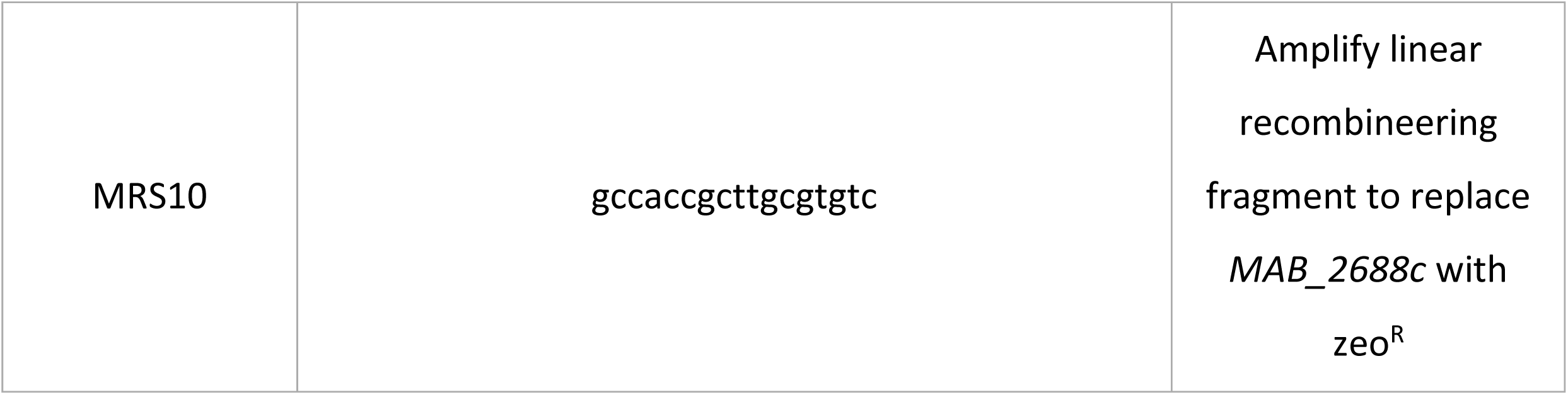
Oligonucleotides used in this study

**Supplemental File 1.** Formulation of tissue culture and mycobacterial media

**Supplemental File 2.** Resampling analysis of relative gene requirements in lung infection model and tissue culture medium versus input library

**Supplemental File 3.** Fatty acid quantitation in various media conditions

